# Modulation of SLP-2 expression protects against alpha-synuclein neuropathology by mitigating mitochondrial dysfunction

**DOI:** 10.1101/2025.06.13.659577

**Authors:** Cyril Bolduc, Maria Paulina Castelo Rueda, Marina Lorente Picón, Giovanna Gentile, Sreehari Kalvakuri, Victoria Soto Linan, Alessandra Zanon, Sofien Laouafa, Martin Lang, Alexandros A. Lavdas, Valentina Gilmozzi, Greta Bernardo, Vincent Coulombe, Charles Gora, Modesto R. Peralta, Véronique Rioux, Claudia Honisch, Peter P. Pramstaller, Paolo Ruzza, Elena Ziviani, Rolf Bodmer, Andrew A. Hicks, Martin Parent, Jorge Soliz, Vincent Joseph, Irene Pichler, Martin Levesque

## Abstract

Parkinson’s Disease (PD) is a progressive neurodegenerative disorder characterized by dopaminergic neuron loss and the accumulation of alpha-synuclein (αSyn)-rich aggregates known as Lewy bodies. Mitochondrial dysfunction is a key contributor to PD pathology, and mitochondrial defects are part of the pathogenic mechanisms induced by αSyn. Stomatin-Like protein 2 (SLP-2) is a mitochondrial scaffold protein that regulates mitochondrial integrity and function. Here, we investigated whether SLP-2 induction can counteract αSyn-induced mitochondrial dysfunction and neurodegeneration. We found that SLP-2 levels were reduced in human PD brains and an A53T αSyn mouse model. Mild overexpression of SLP-2 improved mitochondrial function, reduced oxidative stress, and prevented αSyn-mitochondria interactions in human iPSC-derived neurons. *In vivo*, SLP-2 overexpression protected dopaminergic neurons and motor function, while its depletion exacerbated degeneration and motor deficits in both mouse and *Drosophila* models. These findings suggest SLP-2 as a key regulator of mitochondrial resilience and a potential therapeutic target for PD and alpha-synucleinopathies.

## INTRODUCTION

Parkinson’s Disease (PD) is the second most common and fastest growing neurodegenerative disorder with 6.1 million people living with this disease worldwide (*1*). It affects about 2% of the population over 65 years of age with a prevalence that is globally rising (*2, 3*). The main neuropathological hallmarks of PD include the progressive loss of dopaminergic (DA) neurons in the *substantia nigra pars compacta* (SNc), accompanied by the formation of Lewy bodies, intracellular inclusions composed of aggregated alpha-synuclein (αSyn) as well as other proteins, lipid membranes, and fragments of organelles, including mitochondria (*4, 5*). The subsequent dopamine deficiency in the basal ganglia triggers cellular and synaptic dysfunctions, leading to classic Parkinsonian motor symptoms, which include bradykinesia, rigidity, and tremors (*6, 7*). It remains unclear what triggers the degeneration of DA neurons in PD, but evidence suggests that they are highly susceptible to bioenergetic stress. Midbrain DA neurons have long, highly arborized axonal projections (*8–10*), and manipulating their size increases their bioenergetic burden and susceptibility to mitochondrial stress (*11, 12*).

Most PD cases are classified as idiopathic (∼ 90%), with both genetic and environmental factors playing an important role (*13, 14*). About 10% of cases represent monogenic forms with Mendelian inheritance patterns in genes including *SNCA*, *LRKK2*, *PRKN*, and *PINK1* (*14, 15*). Autosomal dominant single base and genomic multiplication mutations in the *SNCA* gene (e.g. A53T, *SNCA* triplication), encoding the aggregation-prone protein αSyn, cause early-onset PD (*16–18*). Even though past research suggests that αSyn could regulate mitochondrial homeostasis (*19, 20*), accumulating evidence points to mitochondrial defects as part of the pathogenic mechanisms induced by αSyn (*21*). It has been shown that αSyn contains a mitochondrial targeting sequence in its N-terminus (*22*). Additionally, αSyn can interact with TOM20, inhibiting protein import into mitochondria (*23*). This finding aligns with studies reporting reduced levels of mitochondrial import machinery components in both PD and αSyn animal models (*24–26*). Furthermore, αSyn accumulation and oligomerization have been shown to induce mitochondrial fragmentation and dysfunction (*27–32*). Mitochondrial impairment has also been reported in animal models of synucleinopathy (*28, 32–34*). Transgenic mouse lines expressing human A53T mutant αSyn in DA neurons exhibit early-onset mitochondrial abnormalities, characterized by macroautophagy marker-positive cytoplasmic inclusions, which precede neuronal loss (*35, 36*). Impairment of mitochondrial function has also been reported in the DA neurons of mice injected with αSyn pre-formed fibrils, suggesting that mitochondrial dysfunction is a proximal step in the cascade of events leading to neuronal degeneration (*37*). Importantly, the association of αSyn with mitochondria, which is modulated by αSyn dosage and mutation, determines the extent of mitochondrial defects (*19*). Misfolding of αSyn at the mitochondrial membrane is increasingly recognized as an initiating factor for neuronal toxicity. It has been shown that αSyn seeding events in human neurons occur on lipid membrane surfaces, particularly those of mitochondria (*29*). The mitochondrial phospholipid cardiolipin binds A53T αSyn and accelerates its oligomerization, leading to its sequestration within aggregating lipid-protein complexes. This process leads to mitochondrial dysfunction and, ultimately, cell death. These findings further support the notion that mitochondria play a central role in driving αSyn pathology.

Mitochondrial Stomatin-Like Protein 2 (SLP-2, encoded by the *STOML2* gene) is a scaffold protein that binds to cardiolipin and to prohibitins (PHB-1 and PHB-2) and forms microdomains in the inner mitochondrial membrane, which facilitate the assembly of respiratory chain complexes and their function (*38–40*). Cardiolipin synthesis is increased in SLP-2 overexpressing cells, which translates into increased mitochondrial membrane formation and biogenesis (*38*). Moreover, SLP-2 is part of a mitochondrial protease complex (SPY), which regulates the processing of important mitochondrial proteins like PINK1, PGAM5, OPA1, and PRELID1 (*41*). The association of SLP-2 in this complex has an important role in determining the form and function of mitochondria as well as the regulation of cell signaling and survival. We identified SLP-2 as an interaction partner of Parkin, a protein linked to monogenic PD, and found that both SLP-2 and Parkin-depleted cells exhibit similar defects in mitochondrial integrity and bioenergetic function. Importantly, mild induction of SLP-2 expression rescued mitochondrial alterations caused by Parkin deficiency in both human induced pluripotent stem cell (hiPSC)-derived neurons from patients and *Drosophila* PD models (*42*).

We hypothesized that SLP-2, along with its mitochondrial protein and lipid binding partners, could be part of a functional network implicated in αSyn pathology, given that αSyn is known to interact with cardiolipin (*29*). To explore this, we investigated the impact of upregulated SLP-2 expression on mitochondrial function and neuropathology in human neurons and in a mouse model overexpressing A53T αSyn. Our findings reveal that mild enhancement of SLP-2 expression in hiPSC-derived neurons carrying *SNCA* mutations restored mitochondrial function and reduced the mitochondrial localization of αSyn phosphorylated at serine 129 (pS129-αSyn). In the A53T αSyn mouse model, overexpression of SLP-2 in DA neurons conferred neuroprotection against nigrostriatal neurodegeneration, motor deficits, and decline in mitochondrial bioenergetics. Conversely, reduced SLP-2 levels exacerbated DA loss. These findings highlight the potential of SLP-2 as a therapeutic target for mitochondrial dysfunction in PD as well as other synucleinopathies.

## MATERIALS and METHODS

### Human brains

Histological descriptions are based on the analysis of *post-mortem* brains collected from 7 males and 6 females. The mean age and *post-mortem* delays are 76.9 ± 8.9 years and 13.5 ± 6.1 h, respectively (Table S1). The material was obtained from the brain bank of CERVO Brain Research Center. Brain banking and *post-mortem* tissue handling procedures were approved by the Ethics Committee of CIUSSS de la Capitale Nationale. The brains were obtained with written informed consent, and the analyses were performed in conformity with the Code of Ethics of the World Medical Association (Declaration of Helsinki).

The brains were sliced in half along the midline; one side of the brain served for neuropathological examination, while the other side was used for immunohistochemical investigation. The latter hemi-brains were sliced into 2 cm-thick slabs along the coronal plane, and the brainstems were dissected in the mesencephalon. The blocks were fixed by immersion in 4% paraformaldehyde (PFA) at 4°C for 3 days. They were then stored at 4°C in a 0.1 M phosphate-buffered saline (PBS, pH 7.4) solution containing 15% sucrose and 0.1% sodium azide. The slabs containing the *substantia nigra* were cut with a freezing microtome into 50 μm-thick sections that were serially collected in PBS and stored at -20°C in a solution containing glycerol and ethanediol until immunostaining.

### Cell culture

SH-SY5Y-tetON-αSyn cells stably expressing the tet-ON system for tetracycline-inducible expression of *SNCA* were cultured in DMEM/F12 with GlutaMAX-supplement, 10% tetracycline negative FBS (EuroClone), 1% penicillin-streptomycin, 50 µg/ml geneticin and 4 µg/ml blasticidin (Thermo Fisher Scientific). *SNCA* overexpression was induced by the addition of 0.1 µg/ml tetracycline (Sigma-Aldrich) to the cell culture medium. hiPSCs from two control individuals (control 1, hiPSC-802, and control 2, hiPSC-SFC 089-03-07-05A) and two PD patients carrying either a heterozygous triplication of the *SNCA* gene (hiPSC-ND34391) or an A53T mutation (hiPSC-1.1), respectively, were used. hiPSC-802 and hiPSC-1.1 were generated in the laboratory of the Institute for Biomedicine (*43*). hiPSC-SFC 089-03-07-05A was established through the StemBANCC consortium (https://cells.ebisc.org/STBCi033-B/), and hiPSC-ND34391 was obtained by the NINDS Human Cell and Data Repository. hiPSCs were cultured under feeder-free conditions on Matrigel matrix (Corning) in StemMACS iPS-Brew XF (StemMACS) with 1% penicillin-streptomycin (Gibco). hiPSC colonies were passaged enzymatically using 1 mg/ml Collagenase type IV (Gibco) in DMEM/F12 (Gibco). Cell lines are listed in Table S2. The study was approved by the Ethics Committee of the South Tyrolean Health Care System (approval number 102/2014 dated 26.11.2014 with extension dated 19.02.2020).

### Differentiation of hiPSCs into dopaminergic neurons

The direct differentiation of hiPSCs into midbrain DA neurons was conducted as described previously (*44*). Human iPSC colonies were disaggregated into single cells using Accutase (Thermo Fisher Scientific) and replated onto Geltrex-coated (Thermo Fisher Scientific) dishes at a density of 400,000 cells/cm^2^ with Neurobasal/N2/B27 medium (all Thermo Fisher Scientific), containing 2 mM L-glutamine, and supplemented with 10 μM ROCK inhibitor Y-27632 (Miltenyi Biotech), 500 ng/ml recombinant Human Sonic Hedgehog (SHH, R&D System), 0.7 μM CHIR99021 (CH, StemMACS), and SMAD pathway inhibitors SB431542 10 μM (SB, Miltenyi Biotech) and LDN-193189 250 nM (LDN, StemMACS), representing day 0 of differentiation, and cultured from day 1 to 3 without ROCK inhibitor. On days 4 to 9, cells were exposed to 7.5 µM of the Wnt pathway activator molecule CH (day 10, 3 µM). On day 7, LDN, SB and SHH were withdrawn. Upon day 10, maturation of DA neurons was initiated by changing the media to Neurobasal/B27/L-Glu supplemented with 20 ng/ml recombinant Human BDNF (brain-derived neurotrophic factor) (Peprotech), 0.2 mM ascorbic acid (Sigma Aldrich), 1 ng/ml recombinant Human TGF-ß3 (Transforming Growth Factor type 3) (Peprotech), 0.2 mM dibutyryl cyclic-AMP (EnzoLifescience), and 20 ng/ml GDNF (glial cell line-derived neurotrophic factor). On day 12, 10 μM DAPT (Tocris) was added to this midbrain DA differentiation medium. Between days 12 and 15, cells were detached using Accutase and replated at high density (800,000 cells/cm^2^). On day 25 of differentiation, cells were once again replated onto plastic dishes or live imaging chamber slides (ibidi), coated with poly-D-lysine (Sigma Aldrich) and laminin (Sigma Aldrich), for Western blot and imaging experiments. For oxygen consumption rate (OCR) experiments, neurons were replated at day 31, where they were stabilized and further differentiated until day 34.

### Lentivirus production

The human SLP-2 coding sequence was cloned into the lentiviral pER4 vector with phosphoglycerate kinase (PGK) promoter, and virus production was conducted according to standard protocols as we have previously described (*42*). SH-SY5Y-tetON-αSyn cells and hiPSC-derived neurons were infected with a multiplicity of infections of one to three.

### Complex I activity

Mitochondria were isolated from SH-SY5Y-tetON-αSyn cells as previously described (*42*). Briefly, cells were harvested and homogenized in buffer containing 250 mM sucrose, 10 mM Tris, 1 mM EDTA (pH 7.4), and protease and phosphatase inhibitors (Roche Diagnostics). The homogenate was centrifuged twice at 1,500 x g for 10 min. The supernatant containing intact mitochondria was centrifuged at 8,400 x g for 10 min, and the mitochondria-enriched pellet was used for further analysis. Complex I activity was determined by measuring the oxidation of NADH to NAD+ at 340 nm. Citrate synthase was assayed spectrophotometrically according to a published protocol (*45*). The final data were expressed as ratios of complex I/citrate synthase activities.

### ATP synthesis rate

Cellular ATP synthesis rates were measured as previously published (*46*). Briefly, SH-SY5Y-tetON-αSyn cells were harvested, the protein concentration measured by using BCA, and diluted with cell suspension buffer (150 mMol/l KCl, 25 mmol/l Tris-HCl pH 7.6, 2 mmol/l EDTA pH 7.4, 10 mmol/l KPO_4_ pH 7.4, 0.1 mmol/l MgCl_2,_ and 0.1% [w/v] BSA) to a concentration of 1 mg protein per ml. ATP synthesis reaction was initiated by the addition of 250 µl of the cell suspension to 750 µl of substrate buffer (10 mmol/l malate, 10 mmol/l pyruvate, 1 mmol/l ADP, 40 µg/ml digitonin, and 0.15 mmol/l adenosine pentaphosphate). Cells were incubated at 37°C for 10 min. At 0 and 10 min, 50 µl aliquots of the reaction mixture were withdrawn, quenched in 450 µl of boiling 100 mmol/l Tris-HCl, 4 mmol/l EDTA pH 7.75 for 2 min and further diluted 1/10 in the quenching buffer. The quantity of ATP was measured in a luminometer (EnVision 2105 Multimode Plate Reader, Perkin Elmer) with the ATP Bioluminescence Assay Kit (Roche Diagnostics, Basel, Switzerland) following the manufacturer’s instructions.

### Mitochondrial oxygen consumption rate in hiPSC-derived neurons

Mitochondrial oxygen consumption rate (OCR) was measured using the Extracellular O_2_ Consumption Assay Kit (Abcam, ab197243) according to the manufacturer’s instructions. At day 31 of differentiation, 2×10^5^ hiPSC-derived neurons were seeded on a 96-well cell culture plate, previously coated with poly-D-lysine (Sigma Aldrich) and laminin (Sigma Aldrich) and incubated in a CO_2_ incubator at 37°C with NB-B27 medium supplemented with neuronal differentiation factors. Cells recovered for 72 h prior to performing the OCR measurements. For the assay, medium was replaced with 150 µl of fresh NB-B27 medium for routine (basal) OCR measurements, or with 150 µl of NB-B27 medium containing 2.5 µM carbonyl cyanid-p-trifluormethoxyphenylhydrazon (FCCP) for maximal respiration measurements. Next, 10 μl of Extracellular O_2_ consumption reagent were added to each well, except to the blank control, and 100 µl of high-sensitivity mineral oil (pre-heated at 37°C) were added to limit back diffusion of ambient oxygen. Fluorescence intensities were measured using the EnVision 2105 Multimode Plate Reader (Perkin Elmer), pre-heated at 37°C, in 2-minutes-intervals for a total of 200 min, at excitation/emission wavelengths = 355/642 nm. Respiration of cells results in oxygen depletion from the surrounding environment, causing an increase of fluorescence signal. Immediately after performing the Extracellular O_2_ Consumption Assay, the CyQuant™ proliferation assay (Thermo Fisher Scientific) was employed to determine the number of live cells in each well, according to the manufactureŕs instructions. Fluorescence intensity was corrected with the blank control, and OCR was determined by selecting the linear portion of the signal profile (avoiding any initial lag or subsequent plateau) and applying linear regression to determine the slope. The OCR calculated for each well was normalized to the cell number determined by CyQuant fluorescence dye. Fluorescence intensities detecting OCR are expressed as relative fluorescence units (RFU) versus time (min).

### Live Imaging

To assess mitochondrial membrane potential (MMP), mitochondrial superoxide (mROS) production, and mitochondrial morphology in hiPSC-derived neurons under live imaging conditions, 2×10^5^ cells per well were seeded as single cells on an 8-well live imaging chamber slide (ibidi) previously coated with poly-D-lysine (Sigma Aldrich) and laminin (Sigma Aldrich). Neurons were imaged at day 45-50 of differentiation. Cells were incubated with a staining solution containing either 80 nM tetramethylrhodamine methyl ester (TMRM) to assess MMP, 2.5 µM MitoSox Red to evaluate mROS production, or 100 nM Mitotracker green (MTG) to evaluate mitochondrial morphology at 37°C and 5% CO_2_ for 30 min. As a positive control, to represent the background activity of TMRM staining with the concentration used, cells were treated with carbonyl cyanide m-chlorophenylhydrazone (CCCP, 10 µM, 45 min), which completely dissipates the MMP. After washing with pre-warmed HBSS 1X, cells were directly imaged in HBSS at 37°C and 5% CO_2_ using a Leica SP8-X confocal microscope (Leica Microsystems, LAS-X acquisition software), with hybrid detectors and a 63X oil immersion objective. For mean intensity measurements of TMRM and MitoSox Red in thresholded images, Image J software was used (https://imagej.nih.gov/ij/index.html). Mitochondrial fragmentation, assessed based on MTG staining, was calculated using the filament function in Imaris software (version 9.6.0, Bitplane; Oxford Instruments): for each field, the total length of the modelled mitochondrial network was calculated and divided by the number of unconnected parts.

MMP in SH-SY5Y-tetON-αSyn cells was measured by using the cationic dye JC-1 (5,5′,6,6′-tetrachloro-1,1′,3,3′-tetraethylbenzimidazolylcarbocyanine iodide) (5 µg/ml JC-1 in HBSS for 10 min), which exhibits mitochondrial membrane potential-dependent accumulation, indicated by a fluorescence shift from green to red. mROS in SH-SY5Y-tetON-αSyn cells was assessed by MitoSox Red.

### Immunocytochemistry

SH-SY5Y-tetON-αSyn cells and differentiated neurons were grown on 8-well format chamber slides System Nunc™ Lab-Tek™ (Thermo Fisher Scientific), previously coated with poly-D-lysine (Sigma Aldrich) and laminin (Sigma Aldrich) and fixed in a 4% PFA solution. Next, cells were permeabilized in 0.5% Triton X100 - PBS and blocked with blocking solution (3% BSA in PBS) at RT. Immunostaining was performed by addition of primary antibodies against TH (rabbit anti-TH, 1:1,000, Merck Millipore MAB318), pS129-αSyn (mouse anti-pS129-αSyn, 1:200, Abcam ab184674), GRP75 (rabbit anti-GRP75, 1:5,000, Abcam ab53098), and the class III member of the beta tubulin family (mouse anti-TUJ1, 1:1,000, Biolegend). Subsequently, cells were incubated with the appropriate fluorescently labelled secondary antibodies (goat anti-mouse Alexa Fluor 488-conjugated, goat anti-rabbit Alexa Fluor 555-conjugated, Thermo Fisher Scientific), nuclei were stained by adding NucBlue™ Fixed Cell Stain (Thermo Fisher Scientific), and the slides were mounted with Dako Fluorescence mounting medium. Images were acquired using a Leica SP8-X confocal microscope (Leica Microsystems, LAS-X acquisition software), with hybrid detectors and a 63X oil immersion objective.

### Electron microscopy analyses

hiPSC-derived neurons were fixed with 2.5 % glutaraldehyde in 0.1 M sodium cacodylate buffer pH 7.4 ON at 4°C. The samples were postfixed with 1 % osmium tetroxide plus potassium ferrocyanide 1 % in 0.1 M sodium cacodylate buffer for 1 hour at 4°C. After three water washes, samples were dehydrated in a graded ethanol series and embedded in an epoxy resin. Ultrathin sections (60–70 nm) were obtained with an Ultrotome V (LKB) ultramicrotome, counterstained with uranyl acetate and lead citrate, and viewed with a Tecnai G2 (FEI) transmission electron microscope operating at 100 kV. Images were captured with a Veleta (Olympus Soft Imaging System) digital camera and analyzed by ImageJ.

### Western blot analysis

For Western blot analysis, whole-cell homogenates were used. Harvested cells were resuspended in cold RIPA buffer (Thermo Fisher Scientific) supplemented with protease and phosphatase inhibitors (Roche). Protein concentrations were assessed using BCA Protein Assay Kit (Thermo Fisher Scientific), and 10 µg of total protein lysates were loaded per well on a NuPAGE 4-12% Bis-Tris SDS-PAGE gel (Thermo Fisher Scientific). After electrophoresis, proteins were transferred onto a PVDF membrane (BioRad) and probed with antibodies raised in mouse against OPA1 (1:1,000, BD, 612606), SLP-2 (1:1,000, Abcam, ab89025), αSyn (1:1,000, Abnova, MAB5383) and in rabbit against GAPDH (1:5,000, Millipore, MAB374). Subsequently, the blots were incubated with the corresponding HRP-labeled secondary antibodies, and target proteins were detected by enhanced chemiluminescence using Clarity™ ECL Western Kit (BioRad). The chemiluminescence signal was detected using the ChemiDoc™ Touch Imaging System (BioRad) and quantified by densitometry using Image Lab 6.0 analyzer software (BioRad). Optical density values measured for target proteins were normalized by the indicated loading control. For each figure, all blots were processed in parallel and originated from the same experiment.

### Quantitative analysis of RNA expression levels

Total RNA from hiPSC-derived neurons was isolated using the Direct-zol RNA Miniprep Kit (#R2052, Zymo research). RNA integrity was assessed on an Experion Electrophoresis Station (Bio-Rad), and RNA was reverse transcribed using the SuperScript^®^ VILO^TM^ cDNA Synthesis Kit (Thermo Fisher Scientific). qRT-PCR was performed in triplicate on the 96CFX Manager (Bio-Rad) using the Taqman probes Hs00968790_g1 STOML2 FAM-MGB and HS00240906_m1 SNCA FAM-MGB (Thermo Fisher Scientific). The gene expression levels were normalized to the housekeeping gene CYC1 (Hs00357717_m1 CYC1 FAM-MGB) using the ΔΔCt method.

### Drosophila

Flies were raised on standard cornmeal food at 25°C unless otherwise mentioned. To generate dSLP-2 and hSLP-2 transgenic flies, cDNAs of *Drosophila* and human SLP-2 were cloned separately into pUAST-attB vector, and the constructs were injected into attP2 (site-specific integration) flies by BestGene Inc. *SLP-2* (48124^GD^) RNAi was obtained from the VDRC stock center. *TH-Gal4, UAS-StingerGFP, UAS-aSynA53T, elav-Gal4, UAS-LacZ* and *UAS-GFP* RNAi stocks were obtained from Bloomington stock center.

### Negative geotaxis assay

Motor function was assessed as previously described (*42*). Briefly, groups of 10 five-week-old male flies per vial were aged at 25°C and tested for their ability to climb an 8 cm mark within a standard *Drosophila* food vial. The percentage of flies crossing the 8 cm mark within 10 sec was recorded. Each group underwent three trials, and the average pass rate per vial was used for final analysis. A minimum of 100 flies were tested each time in two independent experiments.

### DA neuron counting

Brains were dissected in PBS and fixed in 4% PFA for 30 min followed by three washes in PBS for 15 min each. The images of DA clusters as visualized by TH-Gal4>UAS-StingerGFP were obtained by a Zeiss apotome microscope. The numbers of DA neurons in PPL1 cluster on the posterior side of the brain were determined by the Zeiss imaging software.

### ATP measurement in *Drosophila*

Measurements of ATP were performed as described previously (*47*). Briefly, five heads from 5-weeks old flies were collected and homogenized in 100 µl extraction buffer (100 mM Tris and 4 mM EDTA, pH 7.8) containing 6 M guanidine-HCl, followed by rapid freezing in liquid nitrogen. The samples were boiled for 5 min and cleared by centrifugation at 14,000 x g. Similarly, 10 heads were processed from 2-weeks old flies, and ATP levels were determined by CellTiter-Glo Luminescent Cell Viability Assay (Promega). Relative levels of ATP were determined by dividing the luminescence signal by total protein concentration as measured by BCA.

### Complex I activity in *Drosophila*

Mitochondria-enriched fractions were isolated from whole adult male flies (∼5 weeks old) of the specified genotype (10 flies per sample) as described previously (*47*). Fly heads were gently homogenized in a chilled isolation buffer (250 mM sucrose, 10 mM Tris-HCl pH 7.4, 0.15 mM MgCl₂) using steel beads in a 96-well plate. The homogenate was centrifuged twice at 500 × g for 5 min at 4°C. The resulting supernatant was then centrifuged at 5,000 × g for 5 min at 4°C, and the mitochondrial pellet was resuspended as required. Mitochondrial samples underwent three freeze-thaw cycles in liquid nitrogen before analysis. Complex I activity was measured by monitoring NADH oxidation at 340 nm at 30°C using a Biotek Synergy 2 plate reader. Results are reported as µmol NADH oxidized per minute per citrate synthase activity. Citrate synthase activity was measured by tracking the formation of 5-thio-2-nitrobenzoate at 30°C using the same plate reader. After a 5-min baseline at 412 nm, the reaction was started by adding 0.5 mM oxaloacetate, and the rate was recorded over 15 min.

### Quantification of total αSyn and aggregated αSyn

Fly heads were gently homogenized in 1% NP-40, 25 mM Tris-HCl (pH 7.5), 1.37 M NaCl, 0.027 M KCl supplemented with 1x protease inhibitors (Sigma Aldrich) using steel beads in a 96-well plate. The homogenate was centrifuged at 2,000xg for 15 min at 4°C to remove debris. The resulting clear supernatant was diluted 1:5 using the assay diluent buffer provided with the ELISA kit before measuring total αSyn and αSyn aggregate using ELISA kits from Biolegend (cat# 448607 and (cat# 449407). The αSyn aggregate ELISA kit detects both pathological oligomeric forms and high molecular weight aggregates of αSyn. The ELISA readout was normalized to total brain lysate protein measured by the BCA method.

### Mice

All mouse experiments were performed in accordance with the Canadian Council on Animal Care, and the protocols were approved by the Animal Care Committee of Université Laval. C57BL/6 were purchased from Charles River, and DAT^IRES-Cre^ mice (*48*) and Rosa26^LSL-Cas9^ (*49*) were purchased from Jackson Laboratory (Table S2). These animals were subsequently raised in colonies at the CERVO Brain Research Centre. All animals were housed under controlled temperature (22 ± 2°C) with standard light/dark conditions of 12h. Both males and females were used in all experiments.

### AAV production

The AAV were produced by the Molecular Tools Platform of the CERVO Brain Research Center (https://tools.neurophotonics.ca/). To model PD pathology, adeno-associated virus serotype 2/9 (AAV2/9) encoding the human A53T αSyn expressed under the CMVie-hSyn promoter was generated. For αSyn control AAVs, AAV2/9 encoding mKate under hSyn promoter or AAV2/DJ encoding hSyn-Con/Fon-hChR2(H134R)-EYFP (Addgene, Plasmid #55645) (*50*) were produced. For SLP-2 overexpression experiments, AAV2/9 encoding either hSyn constitutive or Cre conditional double-floxed inverse (DIO) SLP-2-IRES-mCherry were made. As a control for SLP-2 overexpression, AAV2/9 encoding hSyn-DIO-mCherry was produced. For SLP-2 knockdown experiments, double guides RNA (dgRNA) targeting the sequences GAGAGGTTCTCACCGGTTCC and GACTCTGCACATACCGGATT within the exons 2 and 3 of *Stoml2* were encoded into AAV2/9 under the U6 promoter along with CAG-DIO-mCherry-NRN1 fluorescent reporter. As control, AAV2/9 encoding CAG-DIO-mCherry-NRN1 was generated. All AAVs were suspended in 1X PBS with 320 mM NaCl, 5% D-Sorbitol, 0.001% (v/v) Pluronic F-68. AAVs are listed in Table S2.

### Stereotaxic injections

For SLP-2 overexpression cohorts, stereotaxic injections were performed in two to four-month-old DAT^IRES-Cre^ mice (*48*) under isoflurane anesthesia. Head-fixed mice were unilaterally injected in the right side of the SNc with the following coordinates: -3.5 mm (antero-posterior); +1.1 mm (medio-lateral); -4.0 mm (dorso-ventral) from Bregma. A total volume of 1000 nl of AAV solution was injected at a flow rate of 5 nl/sec. AAV2/9 hSyn-αSynA53T and hSyn-Con/Fon-hChr2(H134R)-EYFP were injected at a titer of 5.625×10^12^ genomic copies (GC)/ml, whereas AAV2/9 encoding hSyn-DIO-SLP-2-mCherry and hSyn-DIO-mCherry were injected at a titer of 1.0×10^10^ GC/ml.

Similarly, for high-resolution respirometry experiments, stereotaxic injections were performed in two-to four-month-old male and female C57BL/6 mice under the same surgical conditions and stereotaxic coordinates as described above. In these experiments, AAV2/9 hSyn-mKate was used in place of hSyn-Con/Fon-hChr2(H134R)-EYFP.

For SLP-2 knockdown cohorts, stereotaxic injections were performed in two to four-month-old DAT^IRES-Cre^ crossed with Rosa26^LSL-Cas9^ mice (*49*). Unilateral injection of 1000 nl of AAV at a rate of 5 nl/sec was delivered in the SNc under the same coordinates as the SLP-2 overexpression cohorts. AAV2/9 hSyn-αSynA53T and hSyn-Con/Fon-hChr2(H134R)-EYFP were injected at a titer of 5.625×10^12^ genomic GC/ml, whereas AAV2/9 encoding CAG-DIO-mCherry-NRN1-U6-dgRNA-*Stoml2* and CAG-DIO-mCherry-NRN1 were injected at 2×10^12^ GC/ml.

### Pole test

The pole test was performed 15 weeks post-injection of the AAV. The mice were first acclimatized to the room for one hour and trained one day prior to data collection. Then, the mice were placed face up on top of a 50 cm pole and filmed using a digital camera (GoPro Hero8). The time spent by the mice turning face down (T-Turn) and descending the pole (T-Descend) were measured.

### Cylinder test

The cylinder test was performed 15 weeks post-injection of the AAV. The mice were first acclimatized to the room for one hour before being placed in a transparent glass cylinder 9 cm in diameter and 15.5 cm in height. Then, the mice were filmed over a period of 10 min using a digital camera (GoPro Hero8) prior to data collection. The use of the ipsilateral and contralateral forelimbs to the side of the virus injections was counted as the mice leaned against the cylinder surface.

### Perfusions

Perfusions were performed 16 weeks post-injection of the AAV. Mice were deeply anesthetized using ketamine-xylazine (10 mg/ml; 1 mg/ml) and transcardiacally perfused with 40 ml room-temperature PBS (pH 7.4) followed by 40 ml ice-cold PFA solution (4% PFA in PBS [pH 7.4]). Brains were extracted from perfused animals, fixed overnight in 4% PFA at 4°C, cryoprotected in OCT (Tissue-Tek®, 4583), and stored at -80°C. Samples were then sliced in 60 µm coronal sections with a Leica CM1900 and preserved at -20°C in 30% (v/v) ethylene glycol and 30% (v/v) glycerol buffered to 0.5 M phosphate prior to histologic analysis.

### Immunohistochemistry

Free-floating sections were blocked for 30 min at room temperature with 1% normal donkey serum (NDS) and 0.2% (v/v) Triton X-100 in PBS (pH 7.4). The sections were incubated overnight at 4°C with primary antibodies at optimized concentrations in 1% NDS and 0.2% Triton X-100 blocking solution. The following primary antibodies were used: TH (Pel-Freez Biologicals, P60101, sheep, 1:1,000), NeuN (Millipore, MAB377, mouse, 1:500), SLP-2 (Abcam, ab19884, rabbit, 1:500), mCherry (Novus Biologicals, NBP2-25158, Chicken, 1:1,000; Rockland, 600-401-379, rabbit, 1:1,000), and pS129-αSyn (Wako, 015-25191, mouse, 1:2,000 and 73B3 custom made mouse, 0.76 mg/ml (*51*)). The sections were then washed and incubated for 1 hour at room temperature with secondary antibodies at optimized concentrations in 1% NDS and 0.2% Triton X-100 blocking solution. The following secondary antibodies were used: donkey anti-sheep IgG Alexa Fluor 647 (Jackson Immuno, 713-605-147, donkey, 1:200), donkey anti-sheep IgG Alexa Fluor 488 (Life Technologies, A-11015, 1:400), donkey anti-mouse IgG Alexa Fluor 647 (Life Technologies, A-31571, 1:400), donkey anti-mouse IgG Alexa Fluor 488 (Jackson Immuno, 715-545-150, 1:200), donkey anti-rabbit IgG Alexa Fluor 594 (Jackson Immuno, 711-585-152, 1:200), donkey anti-rabbit IgG Alexa Fluor 647 (Thermo Fisher Scientific, A-31573, 1:400), goat anti-rabbit IgG Alexa Fluor 555 (Life Technologies, A-11011, 1:400), and goat anti-chicken IgG Alexa Fluor 555 (Life Technologies, A-21437, 1:400). The sections were then washed and mounted with Fluoroshield Mounting Medium with DAPI (ab104139) and imaged using a Zeiss LSM700 confocal microscope. Immunofluorescence images were analyzed using ImageJ Software, and the fluorescent signal was measured by averaging the mean pixels intensity value subtracted from the mean pixels intensity value of the background.

### Stereological counting

Six coronal midbrain sections covering the entire SNc with an interval of 120 µm between each slice were selected for stereological counting of DA neurons using TH and NeuN markers. The number of TH+ and NeuN+ cell bodies was estimated by a sampling percentage of 65% using the optical fractionator method of the Stereo Investigator software package (version 11.07; MicroBrightField Biosciences, Williston, VT).

### Oxygen consumption

First, calibration of the Oxygraph 02k-Fluorespirometer (Oroboros Instruments, Innsbruck, Austria) chambers was performed. Mice SNc tissues were manually dissected and weighted between 2-4 mg. The recordings of oxygen consumption were performed at 37°C in a respiration buffer (0.5 mM EGTA, 3 mM MgCl_2_, 60 mM potassium lactobionate, 10 mM KH_2_PO_4_, 20 mM Hepes, 110 mM sucrose, 1 g/l bovine serum albumin). The samples were incubated in the recording chamber for 20 min with 50 μg/ml of saponin as previously described (*52*). For each sample, we have measured mitochondrial oxygen consumption linked to NADH oxidation through the mitochondrial complex I (pyruvate 5 mM, malate 2 mM), mitochondrial oxygen consumption linked to FADH_2_ oxidation through the mitochondrial complex II (succinate 5 mM and rotenone 0.5 μM to block complex I activity) and mitochondrial oxygen consumption linked to NADH-FADH_2_ oxidation through complexes I and II (pyruvate 5 mM, malate 2 mM, succinate 10 mM). The basal oxygen consumption represents the mitochondrial respiratory state 2. After equilibration with the substrates, 2.5 mM ADP was added to the chambers to measure oxygen consumption under normal phosphorylating state 3 (oxphos). We then added 10 μM cytochrome C to assess the integrity of mitochondrial membranes, followed by 2.5 μM oligomycin to block ATP synthesis and measure oxygen consumption due to leakage of protons in non-phosphorylating state 4. Proton gradient uncoupling was obtained by titrating 0.5 μM carbonyl cyanide m-chlorophenylhydrazone (CCCP) until reaching maximum oxygen consumption. We then blocked the activity of complex III with 2.5 μM Antimycin A to measure residual oxygen flux (ROX). ROX was subtracted from all other measurements to report mitochondrial oxygen consumption. To derive oxygen consumption normalized on the weight of brain samples, analysis was performed with the DatLab® software, version 6 (Oroboros Instruments, Innsbruck, Austria).

### *Stoml2* dgRNA validation

Mouse fibroblasts NIH-3T3 (ATCC, CRL-1658) were cultured in Dulbecco’s modified Eagle’s Medium (DMEM, Sigma) supplemented with 10% fetal bovine serum and 1% penicillin-streptomycin (Thermo Fisher Scientific). The cells were co-transfected with Lipofectamine LTX (Invitrogen, 15338-030) with a puromycin-resistant plasmid encoding Streptococcus pyogenes Cas9 (SpCas9) (Addgene, plasmid #62988) (*53*) along with plasmids CAG-DIO-mCherry-NRN1-dgRNA-*Stoml2* or with the control plasmid CAG-DIO-mCherry-NRN1. Three days post-transfection, cells were trypsinized and treated for three days with 2.0 µg/ µl puromycin. The cells were recovered for another two weeks, and genomic DNA was extracted using Monarch® Spin gDNA Extraction Kit (New England Biolabs, T3010). The exons 2 and 3 of *Stoml2* were amplified by PCR using Q5® High-Fidelity DNA Polymerase (New England Biolabs, M0491). The exon 2 of *Stoml2* was amplified using the primers CTCCTATCATGGCCTCAAAGGCTGG and TGGGGATGGGAGGGTGGACTAATAC, whereas the exon 3 was amplified with the primers ACCTTGTAAGGATCCATGATGCG and GGCCTGTCTGCACTTCTAACCTG. The PCR reaction cycle included initial denaturation at 95°C for 4 min, denaturation at 94°C for 30 sec, annealing at 72°C for 45 sec, elongation at 72°C for 1 min, with 30 cycles. The amplicons were sequenced by SANGER at the sequencing platform from CHU of Laval University. The spectrum of nucleotide Insertions and Deletions (INDELS) and the frequency of aberrant base pairs following the expected cut site were measured using Tracking of INDELS by Decomposition (TIDE) (*54*).

### Statistical analyses

Statistical analyses were performed using GraphPad Prism 9. One-way ANOVA was used in experiments comparing more than two groups, followed by Tukey’s *post hoc* test to correct for multiple comparisons. For analyzing differences between two experimental groups, two-tailed unpaired Student’s t-test was utilized. The threshold for significance was set at p<0.05. All experiments were performed in a minimum of three independent biological replicates.

### Sex differences

All males and females were initially analyzed independently for all animal experiments. However, no sexual differences were observed, and the sexes were pooled. In all bar plots, individual data points are annotated by sex using filled dots (males) and empty dots (females), except in two instances (Fig. 9D, E), where sex information was inadvertently not recorded. These data were retained in the analysis due to their scientific value and consistency with the rest of the dataset.

## RESULTS

### SLP-2 levels are reduced in human PD brains and in a mouse model of PD

We first measured SLP-2 levels in human PD donors’ brains. Midbrain samples from *post-mortem* individuals with PD and age-matched controls without a known neurological disorder were selected for histologic analysis (Table S1). Immunofluorescence staining revealed the presence of SLP-2 and pS129-αSyn in TH+ cell bodies of the SNc of both the control and PD human brains (Fig. 1A). As expected, the level of pS129-αSyn was significantly increased in SNc TH+ cell bodies from PD human brains (Fig. 1B). Interestingly, compared to control brains, we observed a significant decrease of SLP-2 levels in the remaining TH+ neurons of PD patient brains (Fig. 1C). Further analysis revealed a near-significant negative correlation between the levels of pS129-αSyn and SLP-2 in TH+ neurons (r^2^=0.2964; p=0.0544, Fig. 1D), suggesting that the accumulation of pathologic αSyn is associated with a reduction of SLP-2 levels in human SNc DA neurons.

**Fig. 1.**
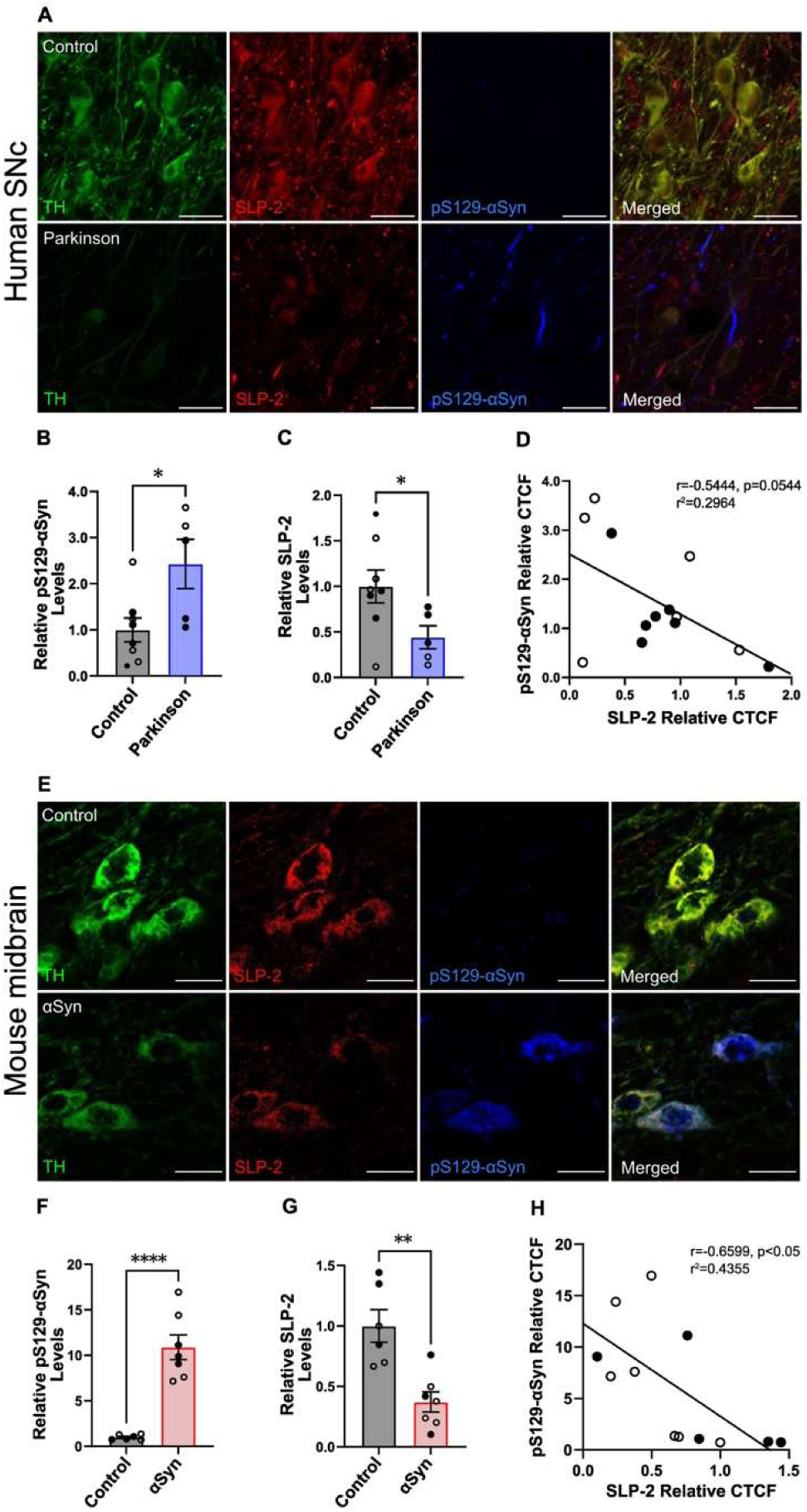
SLP-2 levels are reduced in Parkinsonian human brains and A53T αSyn expression reduces SLP-2 levels in mice. **A)** Representative images of *post-mortem* human brain sections from control donors and patients who suffered from an idiopathic form of PD. *Substantia nigra pars compacta* (SNc) DA neurons were labelled with TH (green), SLP-2 (red) and pS129-αSyn (blue). Scale bar=50 µm. **(B)** Relative quantification compared to the control group of the fluorescent signal of pS129-αSyn and **(C)** of SLP-2 in TH+ neurons. **(D)** Linear correlation between the fluorescence level of pS129-αSyn and SLP-2 in TH+ neurons (linear correlation, r^2^=0.2964, p=0.0544, n=13). **(E)** Representative images of mouse midbrain sections showing SNc DA neurons of two-four months old mice injected in the midbrain with AAV encoding or not A53T αSyn. Animals were sacrificed 16 weeks post-injection of the virus to assess the levels of SLP-2 in the DA neurons of the SNc. Sections were labelled with TH (green), SLP-2 (red) and pS129-αSyn (blue). Scale bar=20 µm. **(F)** and **(G)** Relative quantification compared to the control group of the fluorescent signal of pS129-αSyn and SLP-2 in TH+ neurons. **(H)** Linear correlation between the fluorescence level of pS129-αSyn and SLP-2 in TH+ neurons (linear correlation, r^2^=-0.4355, p<0.05, n=12). Filled circles represent males and open circles represent females. Error bars represent the mean ± SEM. P-values were determined by a two-tailed unpaired Student’s t-test. *p<0.05, **p<0.01 and ****p<0.0001.

To determine whether increased pathological αSyn correlates to decreased SLP-2 levels in SNc DA neurons *in vivo*, we examined this effect in a PD mouse model. We delivered an AAV encoding human A53T αSyn into the SNc of adult mice, with control mice receiving a control AAV. Four months post-injection, the mice were sacrificed for histological analysis. Immunofluorescence staining showed the presence of both SLP-2 and pS129-αSyn in SNc DA cell bodies (Fig. 1E). Quantification of pS129-αSyn fluorescent signal showed a significant increase in pS129-αSyn levels in mice injected with human A53T αSyn compared to controls (Fig. 1F). Notably, we measured a marked reduction in SLP-2 fluorescent signal in the A53T αSyn-injected mice compared to controls (Fig. 1G). Further analysis revealed a significant negative correlation between pS129-αSyn and SLP-2 levels in TH+ neurons (r^2^=0.4355; p<0.05, Fig. 1H), indicating that A53T αSyn induces a reduction of SLP-2 levels in these neurons.

### SLP-2 overexpression rescues mitochondrial malfunction in SH-SY5Y cells with induced αSyn-expression

We have found previously that SLP-2 overexpression rescued mitochondrial alterations in hiPSC-derived neurons from patients with *PRKN* (Parkin) mutations (*42*). Given the observed reduction in SLP-2 levels in SNc DA neurons from PD patients, we investigated whether overexpression of SLP-2 could counterbalance αSyn-induced toxicity on mitochondrial function. To address this, we used inducible human wildtype αSyn-expressing SH-SY5Y cells (SH-SY5Y-tetON-αSyn), in which upregulation of αSyn expression was controlled through the tetracycline-inducible tet-ON system. Quantitative densitometric data from Western blot analyses show that αSyn expression increased significantly (∼10-fold), in comparison to its endogenous expression (uninduced condition) (Fig. S1A, B), and that αSyn overexpression was uniformly distributed in the cell (Fig. S1D). In these cells, we mildly overexpressed SLP-2 by using the pER4-*STOML2* lentiviral vector. Overexpression was confirmed by Western blot analysis, showing a mild (<2-fold), but significant increase in SLP-2 expression (Fig. S1A, C).

Maintenance of the mitochondrial membrane potential ΔΨm is essential for the generation of ATP by the ATP synthase. Therefore, we used ΔΨm as an initial functional readout of mitochondrial health. SH-SY5Y-tetON-αSyn cells showed significantly depolarized mitochondria compared to control cells (p=0.032), a defect that was efficiently restored upon lentiviral SLP-2 transduction (p=0.006) (Fig. 2A, B). Furthermore, the αSyn cells exhibited a functional deficiency of complex I in the electron transport chain, which was restored by SLP-2 expression (p=0.034, Fig. 2C). In parallel, we assessed oxidative stress by measuring superoxide production, which was significantly elevated in the αSyn-expressing cells but was successfully reduced to baseline levels upon SLP-2 overexpression, reaching values comparable to uninduced control cells (p=0.003, Fig. 2D). Additionally, SLP-2 overexpression restored the mitochondrial network integrity, mitigating the fragmentation observed in αSyn-expressing cells (p=0.002, Fig. 2E, F). Collectively, these results highlight the critical role of SLP-2 in preserving mitochondrial structure and function in this cellular model, suggesting its potential as a protective factor against αSyn-induced mitochondrial dysfunction.

**Fig. 2.**
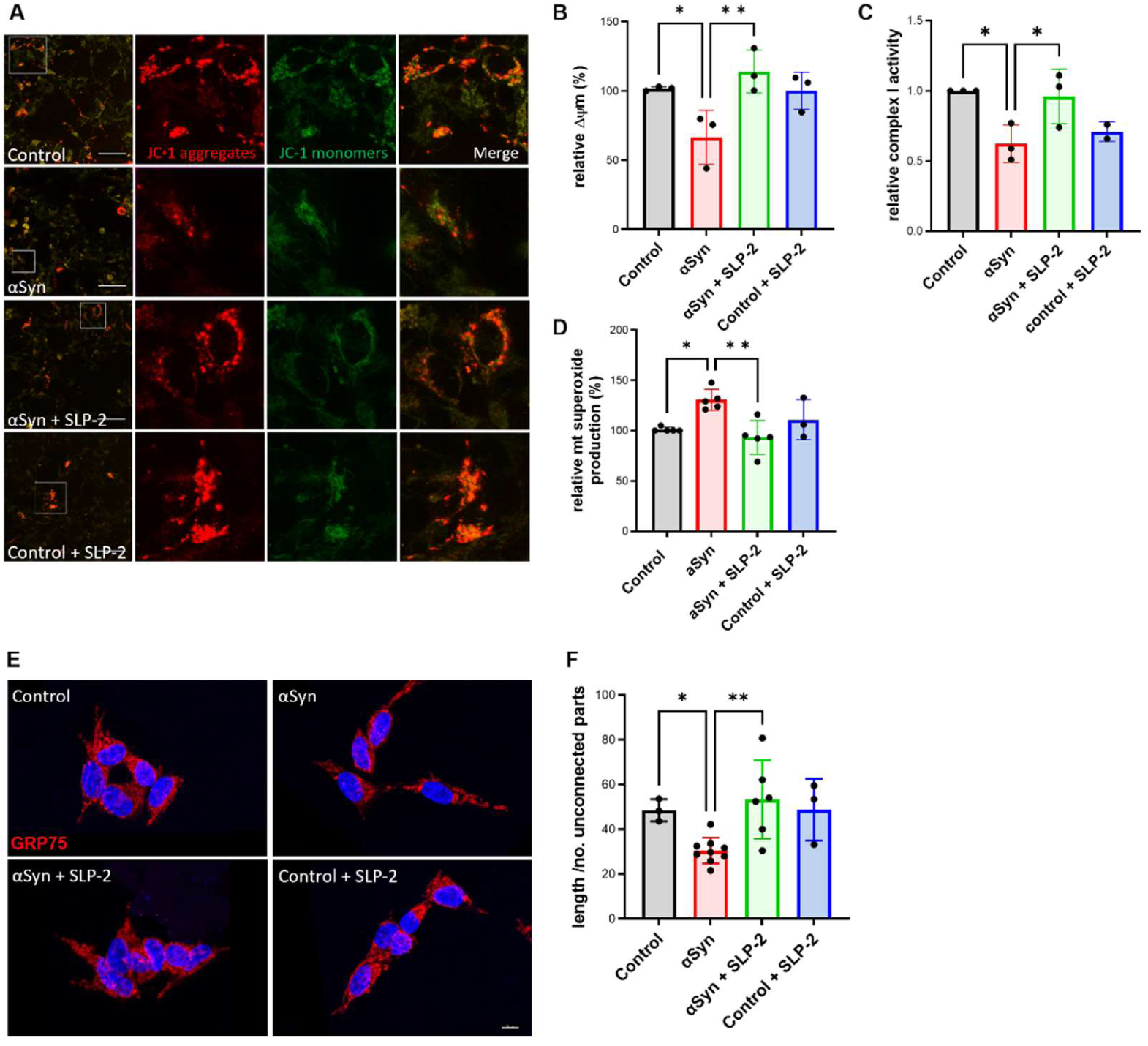
Mitochondrial alterations in SH-SY5Y cells overexpressing αSyn are rescued by moderate SLP-2 expression. **(A)** and **(B)** Upon αSyn overexpression in SH-SY5Y cells, decreased mitochondrial membrane potential, **(C)** reduced complex I activity, **(D)** elevated levels of mitochondrial superoxide, and **(E)** and **(F)** fragmented mitochondrial networks, measured as mitochondrial length over number of unconnected parts, were detected. Moderate, stable increase of SLP-2 expression rescued mitochondrial defects in these cells. n=3 independent experiments. Error bars represent the mean ± SEM. P-values were determined using one-way ANOVA followed by Tukey’s *post hoc* test to correct for multiple comparisons. *p≤0.05, **p≤0.01.

### SLP-2 expression reduces S129 phosphorylated αSyn and its mitochondrial localization in hiPSC-derived neurons with *SNCA* mutations

To validate the mitochondrial effects generated by αSyn toxicity in a disease-relevant preclinical model of PD with endogenous *SNCA* mutations, we assessed mitochondrial phenotypes in two hiPSC-derived DA neuron lines obtained from PD patients, harbouring either an A53T *SNCA* or a 3x*SNCA* mutation (Fig. 3A, Fig. S2), along with two healthy control lines. No difference in endogenous SLP-2 levels was detected in these lines (Fig. S3A-C). *SNCA* mRNA and total αSyn protein levels were increased only in the neurons harbouring the 3x*SNCA* mutation (Fig. S3G-I). By staining for pS129-αSyn, we found significantly increased levels in both the A53T *SNCA* and the 3x*SNCA* lines, compared to the controls (p=0.021; p=0.0003, respectively, Fig. 3B, C). Strikingly, SLP-2 overexpression in these neurons (Fig. S3D-F) reduced pS129-αSyn levels to those comparable to controls, both in neurons with the A53T *SNCA* mutation (p=0.057) and with the *SNCA* triplication (p=0.005) (Fig. 3B, C). Several studies have reported an abnormal association between αSyn and mitochondria (*31, 35, 55*), leading to disruption in various mitochondrial processes (*56*). Consistent with this, we observed an increased co-localization between pS129-αSyn and mitochondria in both mutant lines (p=0.227; p=0.006, Fig. 3B, D). Notably, the overexpression of SLP-2 not only decreased overall levels of pS129-αSyn but also reduced its association with mitochondria (p=0.002, p=0.003, Fig. 3B, D).

**Fig. 3.**
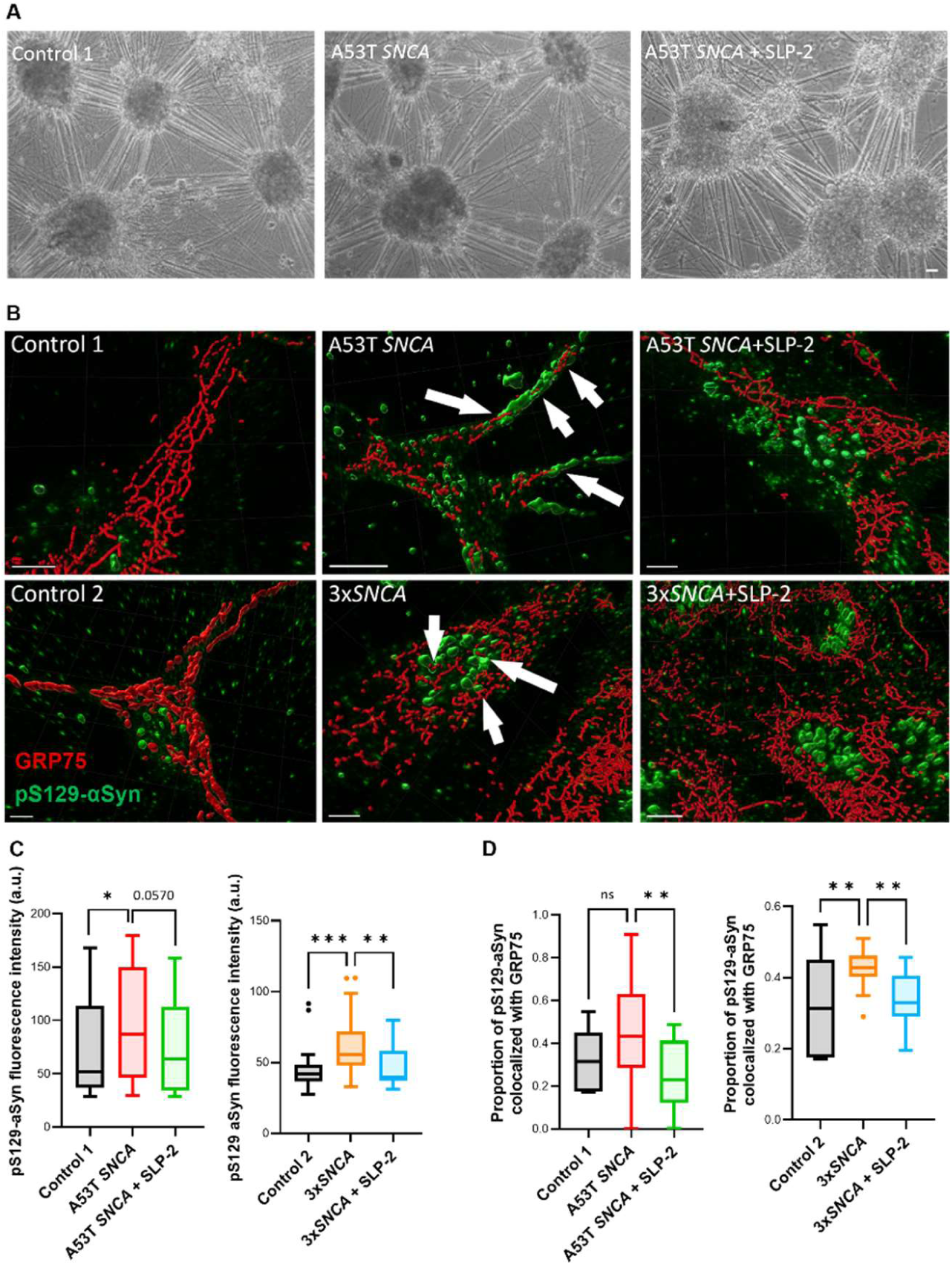
SLP-2 overexpression reduces S129-phosphorylated αSyn in hiPSC-derived neurons with *SNCA* mutations and prevents interaction with mitochondria. **(A)** Representative images visualized by phase contrast inverted microscope of hiPSC-derived neurons at DIV 60. Scale bar=100 µm. **(B)** Representative 3D reconstructions of immunostaining for pS129-αSyn (green) and GRP75 (red) in hiPSC-derived neurons. Scale bar=5 µm. **(C)** Quantification of mean fluorescence intensity of pS129-αSyn as indicator of amount, **(D)** co-localization of pS129-αSyn with GRP75 (mitochondria marker), quantified with the Manders coefficient (proportion of pS129-αSyn in GRP75 signal). n=4-5 independent experiments. Error bars represent the mean ± SEM. P-values were determined using one-way ANOVA followed by Tukey’s *post hoc* test to correct for multiple comparisons. *p≤0.05, **p≤0.01, ***p≤0.001. ns= not significant. a.u.: arbitrary units.

### Bioenergetic impairment in *SNCA* mutant hiPSC-derived neurons is restored by SLP-2

To evaluate mitochondrial function in hiPSC-derived neurons harboring either the A53T mutation or the 3x*SNCA* mutation, we measured oxygen consumption rate (OCR) under basal conditions as well as upon treatment with the protonophore FCCP, leading to mitochondrial uncoupling (maximal respiration) (Fig. 4A). In the mutant neurons, we observed a trend towards reduced basal OCR (Fig. 4B), and challenging these neurons with FCCP to induce maximal respiration revealed impaired oxygen consumption in both mutant lines compared to controls (p=0.026, p=0.007, Fig. 4C). Importantly, upon overexpression of SLP-2 in the patient neurons, oxygen consumption was significantly improved (p=0.006, p=0.009, respectively), reaching levels comparable to those of the control lines (Fig. 4B, 4C).

**Fig. 4.**
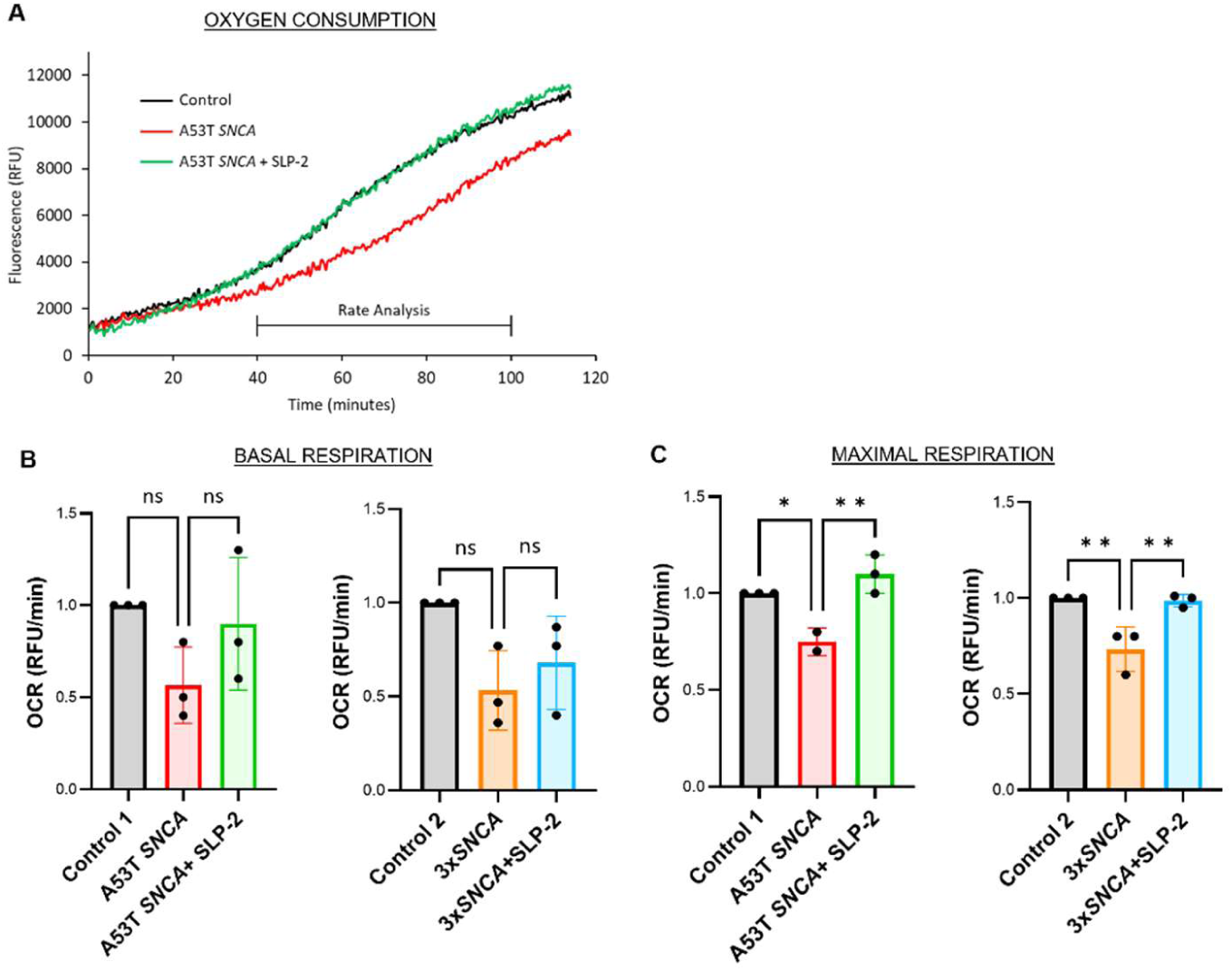
Reduced oxygen consumption in hiPSC-derived neurons with *SNCA* mutations is restored by SLP-2 upregulation. **(A)** Representative maximal respiration curves of the fluorescence signal, generated after the addition of 2.5 µM FCCP. Curves reflect dissolved oxygen in the culture medium of hiPSC-derived neurons of a control line, the A53T *SNCA* mutant line, and the A53T *SNCA* line with SLP-2 overexpression. **(B)** and **(C)** Basal and maximal respiration for the control lines, the *SNCA* mutant lines, and the *SNCA* mutant lines+SLP-2. n=3 independent experiments. Error bars represent the mean ± SEM. P-values were determined using one-way ANOVA followed by Tukey’s *post hoc* test to correct for multiple comparisons. *p≤0.05, **p≤0.01. ns=not significant. RFU: relative fluorescence units.

To assess mitochondrial membrane potential (Δψm) reflecting the electrochemical gradient across the mitochondrial membrane, we utilized the fluorescent dye TMRM and confocal live microscopy (Fig. 5A). In neurons derived from hiPSC carrying the A53T *SNCA* or 3x*SNCA* mutations, we detected significantly depolarized mitochondria, indicating a loss of Δψm (p=0.041, p=0.015, respectively, Fig. 5A, C) compared to the controls. Overexpression of SLP-2 in these mutant neurons restored Δψm to near-normal levels (p=0.063, p=0.046, Fig. 5A, C).

**Fig. 5.**
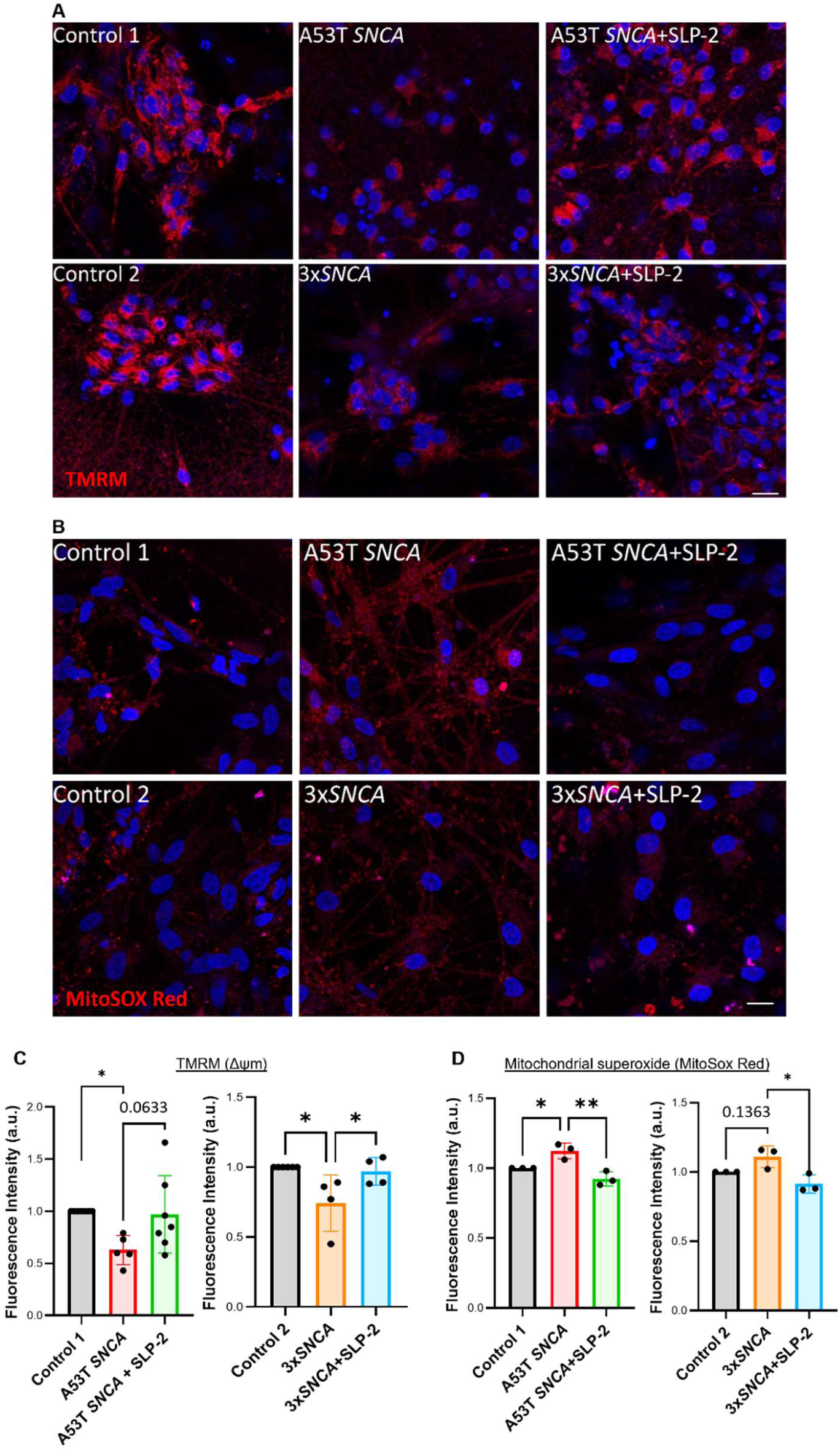
SLP-2 overexpression reinstates mitochondrial membrane potential (Δψm) and reduces oxidative stress in hiPSC-derived neurons with *SNCA* mutations. **(A)** Representative images of TMRM staining in hiPSC-derived neurons at DIV 55. Scale bar=20 µm. **(B)** Representative confocal live imaging of MitoSox Red staining in hiPSC-derived neurons at DIV 55. Scale bar=20 µm. **(C)** Quantification of Δψm as mean fluorescence intensity of TMRM in the control lines, the *SNCA* mutant lines and the *SNCA* mutant lines+SLP-2 overexpression. n=4-5 independent experiments. Error bars represent the mean ± SEM. **(D)** Quantification of mean fluorescence intensity of MitoSox Red in mitochondria of the control lines, the *SNCA* mutant lines, and the *SNCA* mutant lines+SLP-2. n=3 independent experiments. Error bars represent the mean ± SEM. P-values were determined using one-way ANOVA followed by Tukey’s *post hoc* test to correct for multiple comparisons. *p≤0.05, **p≤0.01. a.u.: arbitrary units.

The bioenergetic impairment observed in the patient-derived neurons prompted us to further investigate oxidative stress using the fluorogenic dye MitoSox Red to quantify mitochondrial superoxide (mROS) production. We detected a significant increase in mROS production in the mutant hiPSC-derived neurons (p=0.032, p=0.13, respectively), as indicated by MitoSox-positive mitochondria with an elevated mean fluorescence intensity (Fig. 5B, D), whereas this increased oxidative stress was significantly reduced following SLP-2 expression (p=0.003, p=0.015, Fig. 5B, D). These findings suggest that even a moderate overexpression of SLP-2 expression can have a beneficial impact in the context of αSyn induced pathology in these cellular disease models.

### SLP-2 expression rescues altered mitochondrial shape in hiPSC-derived patient neurons carrying *SNCA*

### mutations

Several studies indicate that mitochondrial fusion and fission events are disrupted in response to increased αSyn expression, leading to mitochondrial network fragmentation in both cellular and animal models (*57, 58*). Consistent with our findings in the SH-SY5Y cell model with induced αSyn expression, mitochondrial network fragmentation was more pronounced in hiPSC-derived neurons carrying *SNCA* mutations compared to control lines (p=0.042, p=0.003, respectively, Fig. 6A), and increasing SLP-2 levels in mutant neurons led to restored mitochondrial structure and reversal of the fragmentation phenotype (p=0.0007, p=0.067, Fig. 6B). To further characterize structural alterations, we performed transmission electron microscopy (TEM) to assess mitochondrial ultrastructure, size and shape (Fig. 6C, D). In neurons carrying the A53T *SNCA* mutation, mitochondria displayed a reduced aspect ratio and a smaller diameter compared to healthy controls (p<0.0001, p<0.0001, respectively), suggesting a shift towards a more rounded morphology and fragmented mitochondrial network (*59*) (Fig. 6C, D, red box). This is further confirmed by the circularity index, which was significantly higher in the mutant line (p<0.0001, Fig. 6C). Overexpression of SLP-2 in mutant neurons mitigated these abnormalities, restoring mitochondrial morphology towards that seen in controls (p<0.0001, p=0.0002, respectively, Fig. 6C).

**Fig. 6.**
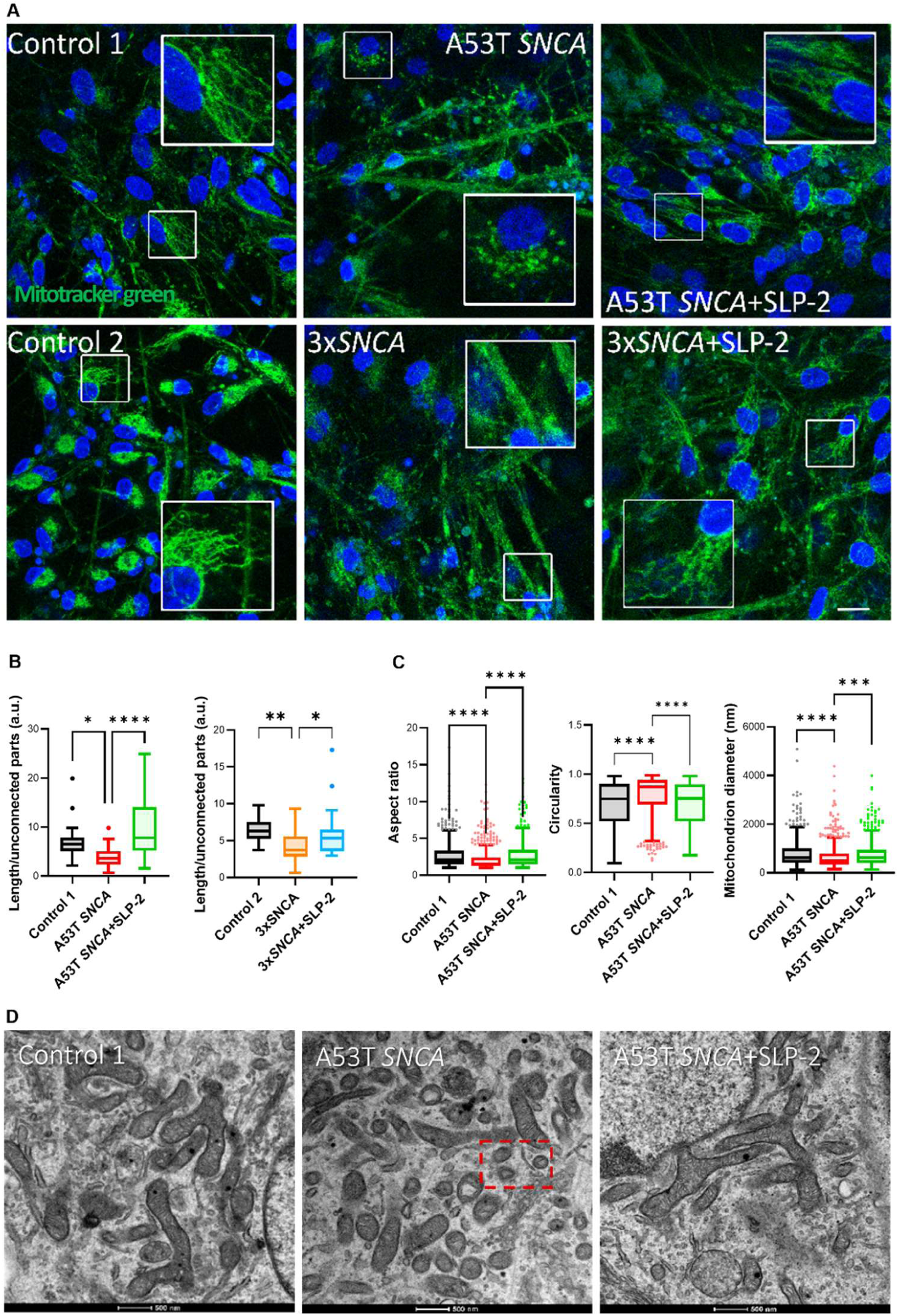
SLP-2 overexpression rescues altered mitochondrial morphology in hiPSC-derived neurons carrying *SNCA* mutations. **(A)** Representative images with Mitotracker green staining in hiPSC-derived neurons at DIV 65. Scale bar=20 µm. **(B)** Mitochondrial morphology was quantified by measuring the total length of mitochondria divided by the number of unconnected parts of the control lines, the *SNCA* mutant lines, and the *SNCA* mutant lines+SLP-2. n=3 independent experiments. **(C, D)** Representative transmission electron micrographs of a control line, the A53T *SNCA* line and the A53T *SNCA* line+SLP-2 and their respective quantification indicating that the mutant line contains smaller mitochondria (aspect ratio) with a decreased average diameter and increased circularity. Accumulation of small round mitochondria is highlighted in the red box. Scale bar=500 nm. Error bars represent the mean ± SEM. P-values were determined using one-way ANOVA followed by Tukey’s *post hoc* test to correct for multiple comparisons. *p≤0.05, **p≤0.01, ***p≤0.001, ****p≤0.0001. a.u.: arbitrary units.

To further investigate these effects, we measured the levels of dynamin-related protein OPA1, a critical mediator of inner mitochondrial membrane fusion and cristae maintenance (*60*) via Western blotting. The levels of the long isoform (L-OPA1) were lower in both mutant lines (A53T mutation, p=0.0421 and 3x*SNCA* mutation, p=0.0427) (Fig. S4A-C), consistent with the observed mitochondrial fragmentation. Overexpression of SLP-2 in mutant neurons, however, did not appear to significantly improve this phenotype (Fig. S4A-C).

### Overexpression of SLP-2 rescues *Drosophila* αSyn A53T overexpression phenotypes

To test whether the beneficial effects of SLP-2 overexpression observed in human neurons translate into whole-organism effects, we analyzed a *Drosophila* model expressing human A53T mutant αSyn. In this model, the knockdown of SLP-2 levels reproduces several phenotypes observed for αSyn A53T OE (Fig. 7). Reduction of SLP-2 combined with αSyn A53T OE led to a further worsening of the αSyn A53T OE phenotype alone. To test whether boosting SLP-2 levels can rescue αSyn A53T OE phenotypes, as we have shown previously in the Parkin RNAi flies (*42*), we produced SLP-2 overexpression transgenes expressing either the *Drosophila* SLP-2 ortholog (dSLP-2) or human SLP-2 (hSLP-2), integrated at the same location in the genome to negate position-dependent changes in transgene expression. Pan-neuronal overexpression of SLP-2 using the elav-Gal4 driver rescued impaired motor function, as indicated by improved climbing pass rates in the negative geotaxis assay (Fig. 7A) and prevented DA neuron loss in A53T mutant flies (Fig. 7B). Additionally, SLP-2 overexpression restored complex I activity and ATP production in αSyn-A53T *Drosophila* PD models (Fig. 7C, D). To determine whether these effects involved modulation of αSyn levels and aggregation, we assessed total αSyn and aggregate formation in brain lysates from *Drosophila* genetic models using an established immunoassay. Notably, SLP-2 overexpression significantly reduced both total αSyn levels and aggregates (Fig. 7E, F). Together, these findings demonstrate that SLP-2 induction restores bioenergetic function and neuronal health in αSyn A53T OE mutant flies.

**Fig. 7.**
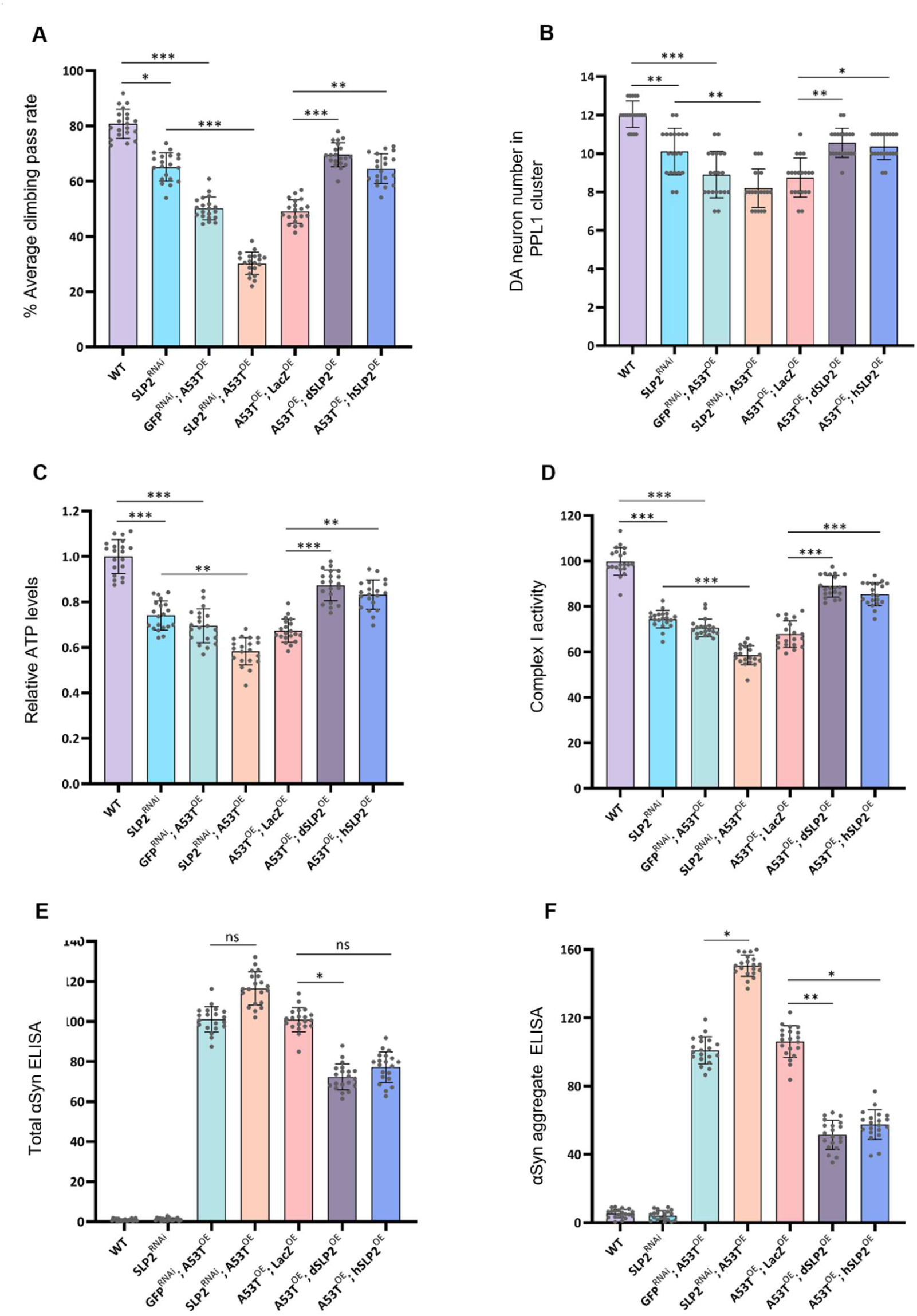
SLP-2 genetically interacts with αSyn and rescues αSyn-A53T mutant phenotypes when overexpressed in the *Drosophila* model. **(A)** Measurement of motor function using negative geotaxis assay. 5-weeks old males aged at 25°C were subjected to climb an 8 cm mark, and the percentage of flies that crossed this mark in 10 seconds was determined. A minimum of 100 flies were tested in repeated trials. *Drosophila* SLP-2 (dSLP-2) or human SLP-2 (hSLP-2), aSyn A53T overexpression transgenes were expressed under the control of *elav*-Gal4. **(B)** Quantification of DA neurons in PPL1 cluster. The numbers of DA neurons visualized by TH>StingerGFP on the posterior side of the brain were counted. **(C)** Measurements of ATP levels from heads of 5-weeks old flies. The relative levels of ATP were determined by dividing the luminescence by the total protein concentration and normalized to wild-type flies. **(D)** complex I activity, normalized to citrate synthase activity, was measured in mitochondria-enriched fractions from 5-weeks old flies. **(E)** and **(F)** Total αSyn levels and αSyn aggregate were measured in brain lysates from 5-weeks old flies using ELISA and normalized to total brain protein. Statistical differences were calculated by one-way ANOVA (non-parametric, Kruskal-Wallis test) followed by Dunn’s *post hoc* test to correct for multiple comparisons. Data is plotted as ± SD. * p ≤ 0.05, ** p ≤ 0.01, *** p ≤ 0.001.

### SLP-2 overexpression prevents motor deficits and protects SNc DA neurons from αSyn toxicity

In our mouse model, A53T αSyn expression led to a 2.5-fold reduction in SLP-2 levels compared to control (Fig. 1E, G). To restore SLP-2 expression to physiological levels, we delivered an AAV encoding a Cre-dependent human SLP-2 gene into the SNc of DAT^IRES-Cre^ mice (*48*). The same viral vector also encoded a Cre-dependent mCherry reporter gene, labelling infected DA neurons (Fig. 8A). Since high-titer injections of the Cre-dependent SLP-2 AAV resulting in ≥four-fold SLP-2 expression were toxic to DA neurons, we optimized the viral dose to 1×10¹⁰ GC/ml. At this concentration, histological analysis confirmed the presence of mCherry+ SNc DA neurons in both control and SLP-2-overexpressing mice (Fig. 8B). Quantification of SLP-2 in mCherry+ neurons confirmed an approximately 1.5-fold increase in expression without signs of toxicity (Fig. 8C).

**Fig. 8.**
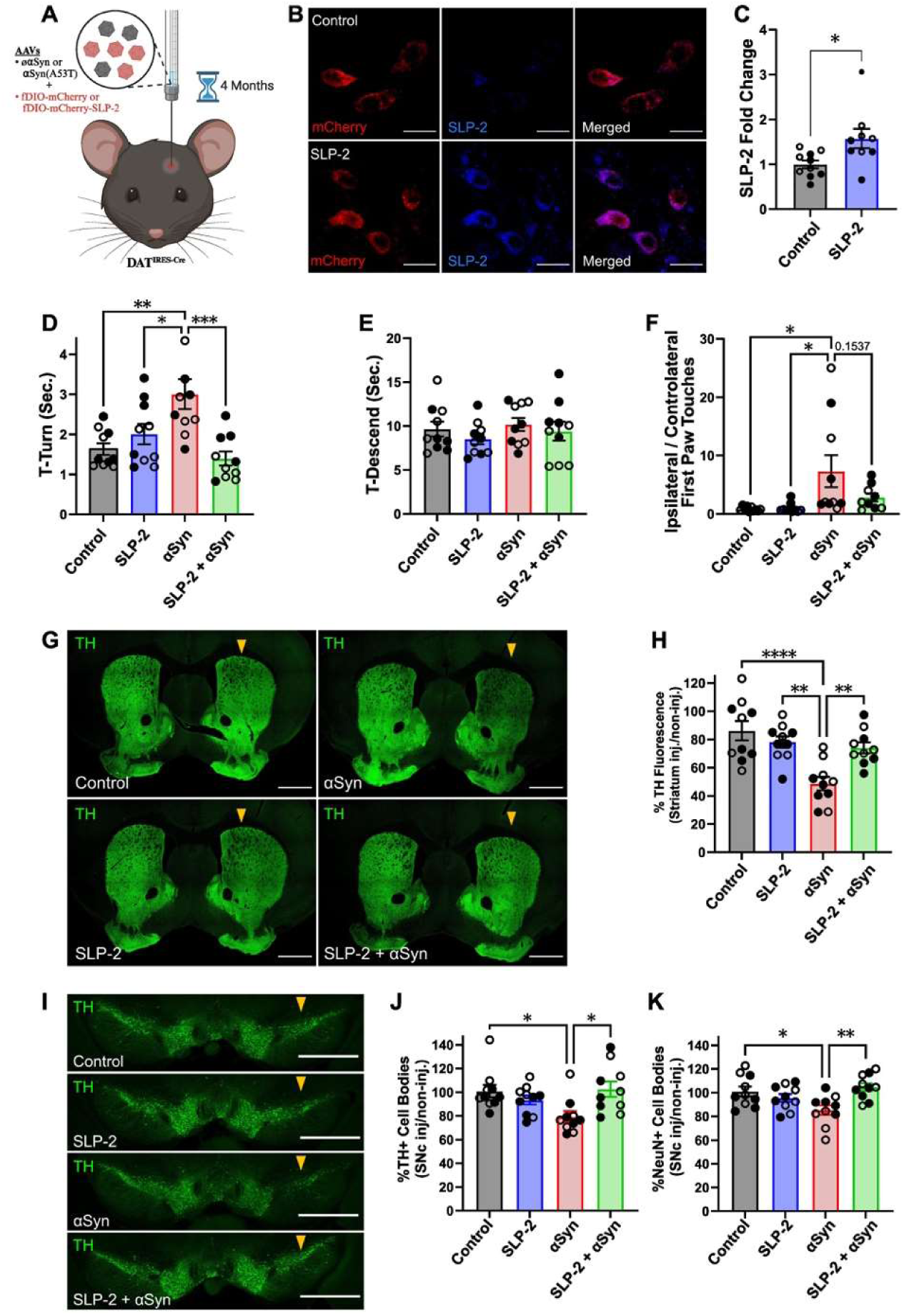
SLP-2 overexpression prevents the appearance of motor deficits and protects SNc DA neurons against degeneration induced by A53T αSyn toxicity. **(A)** Schematic of the experimental procedure. AAV encoding a Cre-dependent copy of the human SLP-2 gene was delivered into the SNc of DAT^IRES-Cre^ mice. The mice were sacrificed 16 weeks post-injection to assess SLP-2 levels in the infected DA neurons, which are marked by the presence of the mCherry reporter gene. **(B)** Representative images of infected DA neurons shown by the mCherry reporter gene (red). SLP-2 (blue) was also labelled. Scale bar=20 µm. **(C)** Relative quantification of the fluorescent signal of SLP-2 in the mCherry+ neurons compared to the control group. Error bars represent the mean ± SEM. P-value was determined by two-tailed unpaired Student’s t-test. **(D)** Quantification of time spent by the mice turning face down (T-Turn) and **(E)** descending the pole (T-Descend). **(F)** Rate of use of the ipsilateral forelimb in the cylinder test. **(G)** Representative images of coronal sections of mouse striatum. DA axonal fibres were labelled with TH (green). The arrows indicate the side of the injection of the AAV. Scale bar=1,000 µm. **(H)** Quantification of the loss of TH fluorescent signal in the ipsilateral dorsal striatum compared to the non-injected side. **(I)** Representative images of mouse SNc coronal sections. The cell bodies of DA neurons were labelled with TH (green). The arrows indicate the side of the injection of the AAV. Scale bar=1,000 µm. **(J)** Quantification of the loss of TH+ and **(K)** NeuN+ cell bodies in the ipsilateral SNc compared to the non-injected side. Filled circles represent males and open circles represent females. Error bars represent the mean ± SEM. P-values were determined by a one-sided ANOVA test followed by Tukey’s *post hoc* test to correct for multiple comparisons. *p<0.05, **p<0.01, ***p<0.001, ****p<0.0001.

**Fig. 9.**
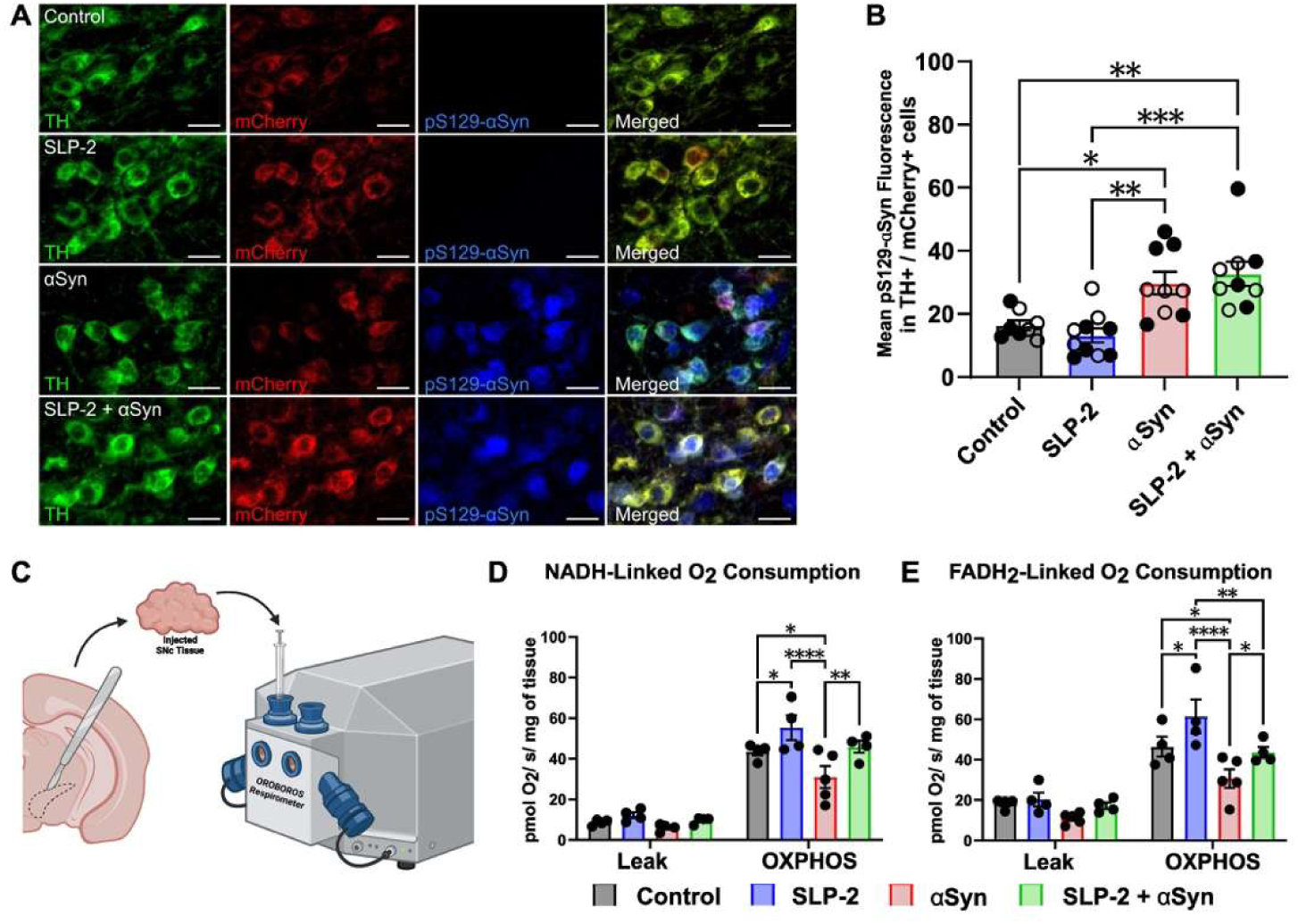
SLP-2 overexpression does not prevent the accumulation of phosphorylated αSyn in mouse midbrain DA neurons but increases mitochondrial oxygen consumption. **(A)** Representative images of mouse SNc coronal sections injected four months prior with AAV encoding either or not SLP-2 and A53T αSyn. The cell bodies of DA neurons were labelled with TH (green), mCherry (red), and pS129-αSyn (blue). Scale bar=20 µm. **(B)** Quantification of the fluorescent signal of pS129-αSyn in the TH+ and mCherry+ neurons. **(C)** Schematic of high-resolution respirometry experimental pipeline. Mice injected for four weeks with AAV encoding or not SLP-2 and A53T αSyn were dissected to extract the SNc tissue and measure the NADH and FADH_2_-linked mitochondrial oxygen consumption. **(D)** NADH-linked and **(E)** FADH_2_-linked oxygen consumption during the leak and OXPHOS phases. Filled circles represent males and open circles represent females. Error bars represent the mean ± SEM. P-values were determined by a one-sided ANOVA test followed by Tukey’s *post hoc* test to correct for multiple comparisons. *p<0.05, **p<0.01, ***p<0.001, ****p<0.0001.

Next, we tested whether SLP-2 overexpression conferred neuroprotection against αSyn toxicity in our PD mouse model. Two to four-month-old DAT^IRES-Cre^ mice received a unilateral injection of either a control AAV or an AAV overexpressing SLP-2 in DA neurons. Simultaneously, a second AAV encoding either A53T αSyn or a control sequence was co-injected. Motor performance was assessed 15 weeks post-injection using the pole and cylinder tests. In the pole test, A53T αSyn-expressing mice showed prolonged turn times compared to controls, whereas mice overexpressing SLP-2 did not exhibit this deficit (Fig. 8D). No significant difference was observed in descent time across groups (Fig. 8E). In the cylinder test, A53T αSyn-expressing mice exhibited a preference for the ipsilateral forelimb, suggesting motor impairment, while SLP-2 overexpression appeared to mitigate this effect, though the difference did not reach statistical significance (p = 0.1537) (Fig. 8F). Mice were sacrificed 16 weeks post-injection for histological analysis. Quantification of TH+ fluorescent signal in the dorsal striatum showed that A53T αSyn induced significant DA axonal denervation, which was prevented by SLP-2 overexpression (Fig. 8G, H). In the SNc, stereological counts of TH+ and NeuN+ neurons confirmed DA neuron loss in A53T αSyn-expressing mice, which was also prevented by SLP-2 overexpression (Fig. 8I-K). Together, these results indicate that SLP-2 overexpression prevents motor deficits and protects SNc DA neurons against degeneration caused by A53T αSyn toxicity.

### SLP-2 overexpression ameliorates early bioenergetic deficits but does not prevent the accumulation of phosphorylated αSyn in mouse midbrain DA neurons

Since our *in vitro* characterization showed that mild SLP-2 overexpression reduces phosphorylated αSyn accumulation in hiPSC-derived DA neurons carrying *SNCA* mutations, we assessed whether a similar effect occurs following viral-mediated expression of A53T αSyn *in vivo*. To this end, we stained midbrain sections for pS129-αSyn (Fig. 9A). Quantification of fluorescent intensity in infected DA neurons confirmed that A53T αSyn expression triggers pS129-αSyn accumulation. However, in contrast to our hiPSC-derived neuron model, SLP-2 overexpression did not prevent this accumulation *in vivo* (Fig. 9B).

To investigate how SLP-2 mitigates αSyn-induced DA neuron degeneration, we examined whether SLP-2 could mitigate mitochondrial dysfunction in our PD mouse model independently of phosphorylated αSyn accumulation. Mitochondrial oxygen consumption was measured in dissected SNc by high-resolution respirometry as previously described (*61–63*). Given that four months of A53T αSyn expression results in significant DA neuron loss (Fig. 8I-K), changes in oxygen consumption at this stage could reflect differences in cell number rather than mitochondrial deficits *per se*. To control for this, we injected the SNc of C57BL/6 mice with AAV2/9 encoding either A53T αSyn or mKate as a control, both expressed under the hSyn promoter. In parallel, mice were co-injected with another AAV2/9 encoding either SLP-2 along with a mCherry reporter or mKate alone. To assess mitochondrial function at early disease stages, we performed high-resolution respirometry four weeks post-injection on dissected SNc tissues (Fig. 9C). No significant difference was measured in the leak state for complex I and complex II (Fig. 9D-E), indicating comparable proton leakage, membrane fluidity, and uncoupling protein activity across groups. However, when measuring the mitochondrial respiratory capacity in the ADP-stimulated OXPHOS state, we found a dual regulatory effect between A53T αSyn and SLP-2 on complex I and complex II activity. A53T αSyn expression reduced NADH and FADH_2_-linked oxygen consumption compared to controls, but SLP-2 overexpression rescued this deficit. Interestingly, SLP-2 overexpression alone enhanced NADH and FADH_2_-linked oxygen consumption rates beyond control levels (Fig. 9D-E). Altogether, these results indicate that A53T αSyn expression induces early bioenergetic deficits *in vivo*, which can be rescued by SLP-2 overexpression independently of pathological phosphorylated αSyn accumulation.

### SLP-2 knockdown in midbrain DA neurons exacerbates αSyn-mediated toxicity

To investigate whether reducing SLP-2 levels in mouse midbrain DA neurons affects αSyn-mediated toxicity, we first designed a dgRNA targeting the introns 2 and 3 of the *Stoml2* gene (Fig. S5A). These dgRNA were encoded into a plasmid and co-transfected into mouse NIH-3T3 cells along with a plasmid encoding SpCas9 and a puromycin resistance gene (*53*). After three days of selection with puromycin, cells were recovered for two weeks and genomic DNA was extracted. PCR amplification and sequencing of introns 2 and 3, followed by TIDE analysis, confirmed that the dgRNA successfully induced double-stranded breaks at the expected cut sites, leading to frameshift mutations in at least 62% and 57% of the sequenced copies, respectively (Fig. S5B, C). The validated dgRNA was then packaged into an AAV vector. To achieve SLP-2 reduction in midbrain DA neurons, we injected this AAV into the SNc of DAT^IRES-Cre^ mice crossed with Rosa26^LSL-Cas9^ (*49*), which express SpCas9 specifically in DA neurons. Given that most mitochondrial proteins have a long half-life, lasting as long as 40 days (*64, 65*), we collected the brains four months post-injection to ensure sufficient SLP-2 depletion. Immunohistochemical analysis confirmed a 50% reduction in SLP-2 levels within TH+ neurons (Fig. 10A, B).

**Fig. 10.**
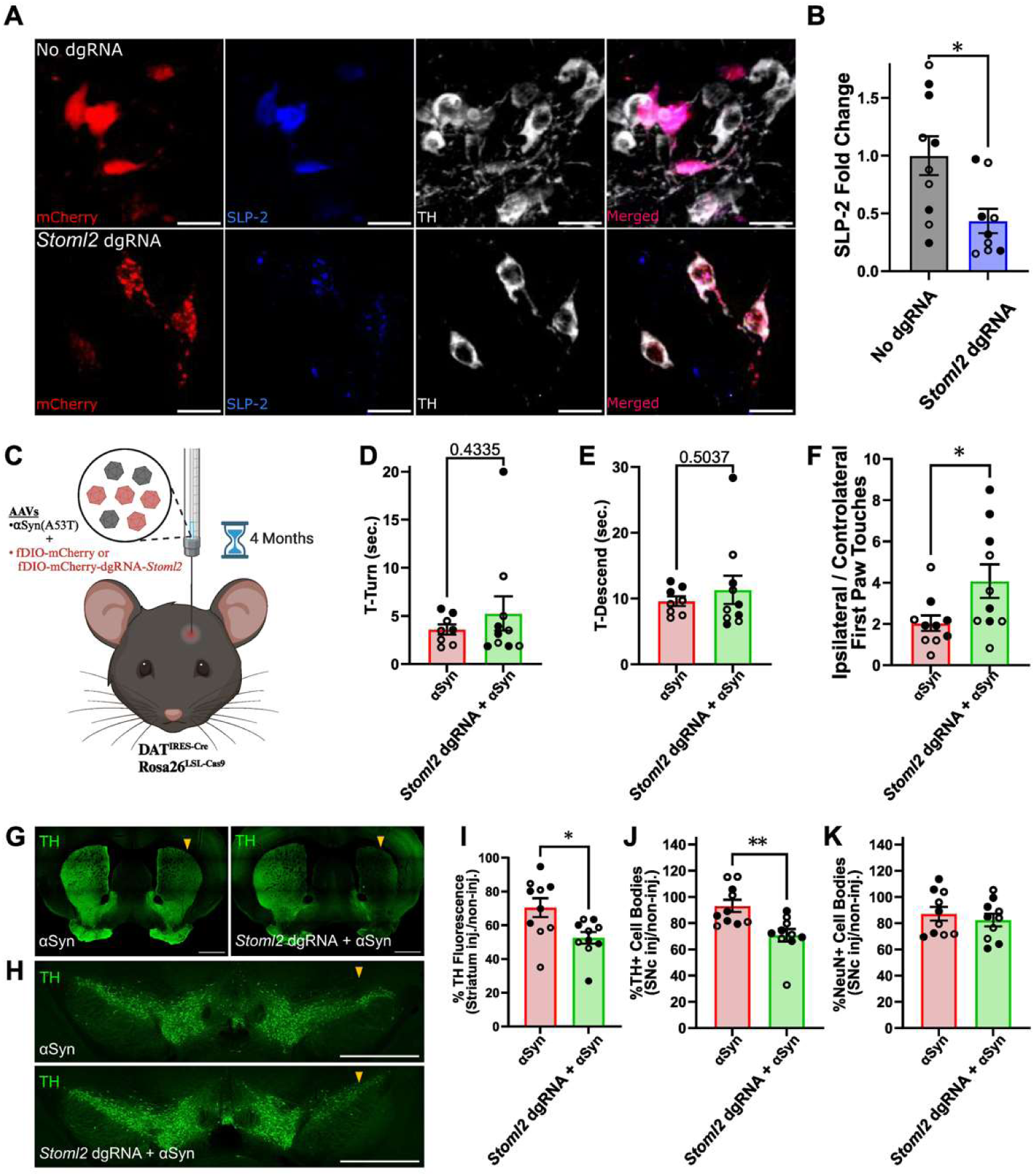
SLP-2 reduction in midbrain DA neurons potentiates αSyn-mediated reduction of TH levels in both the striatum and SNc while leaving the DA cell bodies intact. **(A)** Representative images of infected DA neurons from midbrain sections of DAT^IRES-Cre^Rosa26^LSL-Cas9^ mice injected for four months with AAV encoding a DIO-mCherry fluorescent reporter with or without dgRNA against *Stoml-2*. Filled circles represent males, and open circles represent females. Infected DA neurons were labelled by the mCherry reporter gene (red), with SLP-2 (blue) and TH (green). Scale bar = 20 µm. **(B)** Relative quantification of the fluorescent signal of SLP-2 in the mCherry+ neurons relative to the control group. P-value was determined by two-tailed unpaired Student’s t-test. **(C)** Schematic of the experimental pipeline to assess how SLP-2 reduction is catalyzing A53T αSyn-mediated toxicity. DAT^IRES-Cre^Rosa26^LSL-Cas9^ mice were injected with A53T αSyn encoding AAV along with another AAV encoding or not the dgRNAs against *Stoml-2*. The mice were kept for four months post-injection to conduct motor assessments and histologic analysis. **(D)** Quantification of time spent by the mice turning face down (T-Turn) and **(E)** descending the pole (T-Descend). **(F)** Rate of use of the ipsilateral forelimb in the cylinder test. **(G)** Representative images of coronal sections of mouse striatum. DA axonal fibres were labelled with TH (green). The arrows indicate the side of the injection of the AAV. Scale bar = 1,000 µm. **(H)** Representative images of mouse SNc coronal sections. The cell bodies of DA neurons were labelled with TH (green). The arrows indicate the side of the injection of the AAV. Scale bar = 1,000 µm. **(I)** Quantification of the loss of TH fluorescent signal in the ipsilateral dorsal striatum compared to the non-injected side. **(J)** Quantification of the loss of TH+ cell bodies in the ipsilateral SNc compared to the non-injected side. **(K)** NeuN+ cell bodies in the ipsilateral SNc compared to the non-injected side. Error bars represent the mean ± SEM. P-values were determined by a one-sided ANOVA test followed by Tukey’s *post hoc* test to correct for multiple comparisons. *p<0.05, **p<0.01.

To determine whether SLP-2 reduction enhances αSyn toxicity, we unilaterally injected DAT^IRES-Cre^Rosa26^LSL-Cas9^ mice with AAV encoding A53T αSyn, along with either AAV encoding *Stoml2* dgRNA and a Cre-dependent mCherry cassette or a control AAV encoding only the Cre-dependent mCherry cassette (Fig. 10C). Motor assessment conducted 15 weeks post injection revealed no significant differences in the pole test (Fig. 10D, E). However, the cylinder test showed a significant increase in ipsilateral forelimb use in mice injected with *Stoml2* dgRNA (Fig. 10F). Histological analysis performed the week after revealed that SLP-2 knockdown exacerbates αSyn-mediated toxicity in DA neurons. In mice injected with AAV encoding *Stoml2* dgRNA, we observed a greater loss of TH+ striatal fibers and a more pronounced reduction in TH+ cell bodies in the SNc compared to the contralateral, non-injected side (Fig. 10G–J). This reduction in TH is likely indicative of a cellular dysfunction rather than overt neuronal loss. Supporting this, numerous mCherry+ neurons in the SNc lacked TH expression (Fig. S6), suggesting that DA neurons remain present but undergo a loss of their somatodendritic phenotype. Consistently, NeuN+ cell counts in the SNc were not significantly decreased (Fig. 10H, K), further indicating that the observed reduction in TH immunoreactivity reflects a downregulation of TH expression rather than DA neuron degeneration. Of note, the genetic background impacted vulnerability to αSyn toxicity, as AAV-A53T αSyn induced a stronger phenotype in DAT^IRES-Cre^ mice (Fig. 8G-K) than in DAT^IRES-Cre^Rosa26^LSL-Cas9^ mice (Fig. 10G, I), independently of SLP-2 levels.

## DISCUSSION

In this study, we used complementary preclinical *in vitro* and *in vivo* models to investigate molecular mechanisms of αSyn-induced mitochondrial pathology and neurodegeneration, focusing on the role of the mitochondrial scaffold protein SLP-2 in mitigating these effects. Previously, we have found that moderate SLP-2 expression ameliorates mitochondrial defects caused by reduced Parkin function in SH-SY5Y neuroblastoma cells and hiPSC-derived neurons from PD patients with *PRKN* (Parkin) mutations (*42*). The protective effect of SLP-2 was also replicated in a Parkin-deficient *Drosophila* model, where DA neurons loss and the associated motor dysfunction were restored (*42, 66*). Here, we provide evidence that an enhancement of SLP-2 expression mitigates mitochondrial dysfunction in hiPSC-derived patient neurons with *SNCA* mutations as well as *Drosophila* and mouse models overexpressing A53T αSyn. Furthermore, SLP-2 overexpression protected against motor dysfunction and DA neuron degeneration in both the A53T αSyn fly model and in mice following A53T-αSyn injection in the SNc.

In midbrain neurons of the mouse overexpressing A53T αSyn, SLP-2 protein expression was significantly reduced, mirroring findings in SNc DA neurons from human *post-mortem* PD brains, where SLP-2 levels negatively correlated with pS129-αSyn levels. These results align with previous studies suggesting that αSyn limits mitochondrial protein import (*23–26*). However, this reduction in SLP-2 levels was not observed in hiPSC-derived neurons carrying the endogenous A53T mutation, possibly due to the lower expression levels of the mutant protein. To directly test the effect of SLP-2 deficiency in neurons, we used CRISPR/Cas9 to reduce SLP-2 levels in DA neurons of the A53T αSyn mice, which exacerbated motor deficits and reduction of TH in the cell bodies and striatal fibres. Similarly, in *Drosophila*, reducing SLP-2 levels exacerbated αSyn-mutant phenotypes, supporting a genetic interaction between SLP-2 and αSyn. These findings suggest that residual SLP-2 may confer partial protection in both αSyn mutant mice and the αSyn fly model. Given the critical role of SLP-2 in mitochondrial membrane organization, respiration, and biogenesis via cardiolipin interactions (*38, 40, 41*), we tested SLP-2 overexpression as a potential therapeutic approach. SLP-2 and its mitochondrial protein and lipid binding partners may form an interactive network affected by αSyn in PD (*29*).

Consistent with prior studies, where αSyn accumulation and oligomerization in *SNCA* mutant hiPSC-derived neurons were associated with impaired mitochondrial function (*29, 55, 67*), we detected reduced oxygen consumption, which is an important indicator of OXPHOS activity, mitochondrial membrane depolarization, and increased mROS formation. SLP-2 overexpression effectively restored these bioenergetic defects. The impact of A53T-mutant αSyn overexpression on mitochondrial function was also assessed in *Drosophila* and mouse SNc tissue, replicating the compensatory effect of SLP-2 overexpression *in vivo*. In line with these data, downregulation of SLP-2 in HeLa cells, mouse embryonic fibroblasts, and T-cells from T cell-selective SLP-2 knockout mice has been shown previously to result in defective mitochondrial respiration and respiratory chain supercomplex formation (*39, 40, 68*), while upregulation of SLP-2 in T-cells improved respiratory phenotypes (*38*).

Localization of pS129-αSyn to mitochondria has been observed in neuronal and mouse models of PD, as well as *post-mortem* brain samples of patients with PD and other synucleinopathies (*31, 35, 55*). Misfolded αSyn at the mitochondrial membrane was found to initiate neuronal toxicity, as αSyn seeding events occur on lipid membrane surfaces and in particular on mitochondrial membranes (*29*). We found increased levels and mitochondrial localization of pS129-αSyn in *SNCA* mutant hiPSC-derived neurons, which was attenuated by SLP-2 overexpression. In brain lysates from *Drosophila*, SLP-2 overexpression significantly reduced both total αSyn levels and aggregates. Conversely, SLP-2 overexpression did not alter pS129-αSyn levels in TH+ SNc neurons of A53T αSyn mice, likely because AAV-mediated αSyn overexpression in this model is much higher than the endogenous levels present in *SNCA* mutant hiPSC-derived neurons. Furthermore, we did find that mild overexpression of SLP-2 is neuroprotective but could be toxic for DA neurons if expressed higher than four times its endogenous level, holding implications for potential interventional or therapeutic strategies that target SLP-2. Finally, in line with our results, robust degeneration induced by injection of AAV expressing A53T αSyn has been reported in DAT^IRES-Cre^ (*69*)^69^ but we failed to replicate it in another transgenic background (Rosa26^LSL-Cas9^ mice), emphasizing that different genotypes could show different levels of vulnerability to αSyn toxicity.

Several studies have highlighted a specific interaction between αSyn and the mitochondrial phospholipid cardiolipin (*57, 70, 71*). Cardiolipin is primarily located in the inner mitochondrial membrane, where it constitutes 15-20% of the total lipid content (*72*) and at inner-outer membrane contact sites (*73*). Using a single-molecule Förster resonance energy transfer (smFRET) biosensor, it was shown that exogenously added A53T-αSyn monomers convert into two distinct oligomeric states in primary rodent neurons. In control hiPSC-derived cortical neurons, externally applied A53T-αSyn monomers co-localized with endogenous cardiolipin (visualized by NAO staining), and neurons treated with wildtype αSyn oligomers exhibited increased aggregate formation in A53T mutant cells compared to controls and increased co-localization of these aggregates with cardiolipin, suggesting that cardiolipin promotes the aggregation of αSyn and becomes incorporated into the resulting aggregates. Furthermore, hiPSC-derived A53T mutant neurons exhibited increased endogenous αSyn aggregation, and greater uptake of exogenously added A53T αSyn. These changes were associated with bioenergetic defects, mitochondrial permeability transition pore opening and cell death (*29*). Notably, increased mROS levels after αSyn oligomer exposure can induce local mitochondrial lipid and protein peroxidation, while inhibiting oxidation prevented mitochondrial dysfunction (*35*).

Cardiolipin-enriched microdomains at the inner mitochondrial membrane also contain SLP-2, which binds cardiolipin (*38, 68*) and interacts with membrane-organizing proteins, including prohibitins, and several MICOS (mitochondrial contact site and cristae organizing system) components (*38, 74*). Enzymes involved in cardiolipin synthesis form a large mitochondrial complex that associates with these cardiolipin-binding proteins, suggesting that cardiolipin synthesis is linked to mitochondrial membrane organization (*75*). These previous data raise the hypothesis that a tight regulation of SLP-2 levels might be crucial in preventing or modulating contacts between mutant αSyn and cardiolipin resulting in improved mitochondrial membrane organization and respiratory chain function. In line with this, we observed reduced colocalization of pS129-αSyn with mitochondria in SLP-2 overexpressing hiPSC-derived neurons compared to the αSyn mutant lines lacking exogenous SLP-2 expression.

A previous study has also reported that hiPSC-derived A53T and E46K mutant neurons display highly fragmented mitochondria, reduced mitochondrial membrane potential and increased mitophagy. Mitochondrial fragmentation was preceded by cardiolipin translocation to the outer mitochondrial membrane in both mutant neurons and A53T transgenic mice (*55*). In our study, confocal and electron microscopy analysis confirmed mitochondrial fragmentation in A53T and 3x*SNCA* mutant hiPSC-derived neurons. SLP-2 expression restored mitochondrial morphology, supporting its role in mitochondrial fusion (*76*). Furthermore, L-OPA1 levels, which were significantly decreased in the mutant neurons, were slightly increased by SLP-2 expression, consistent with its function in association with the SPY complex (*41*). Overall, SLP-2 plays an important role in the mitochondrial homeostasis, particularly during oxidative stress, by stabilizing mitochondrial membrane proteins and promoting cardiolipin biosynthesis (*38, 41, 68*). Recently, we identified the *C. elegans* ortholog of SLP-2 (STL-1) as a key regulator of the mitochondrial unfolded protein response (mtUPR), a stress response that maintains mitochondrial proteostasis. STL-1/SLP-2 was shown to protect against exogenous oxidative stress by directly interacting with phosphatidic acid, a cardiolipin precursor, thereby mediating mtUPR (*77*). Importantly, genes linked to SLP-2/cardiolipin pathway, such as Cardiolipin synthase 1 (*CRLS1*) and Fatty Acid Synthase (*FASN*), were recently associated with PD risk in PD genome-wide association studies (*78, 79*), reinforcing the involvement of this mitochondrial pathway in PD. Reduced SLP-2 levels in *post mortem* PD brains further support its role in PD pathology, potentially through modulation of αSyn-induced mitochondrial dysfunction.

In conclusion, our findings demonstrate that SLP-2 mitigates αSyn-induced mitochondrial dysfunction and neuropathology. AAV-mediated overexpression of SLP-2 improved motor performance, protected DA neurons, and preserved striatal DA terminals in the A53T-αSyn mouse model. Consistent with these results, SLP-2 restored mitochondrial integrity and reduced pS129-αSyn levels and αSyn aggregates in *Drosophila* and patient-derived hiPSC-neurons carrying *SNCA* mutations. Combined with prior evidence of its protective effects in *PRKN* (Parkin)-mutant cellular and *Drosophila* PD models (*42*), our data highlight SLP-2 as a promising target for enhancing mitochondrial function and neuronal survival in PD subtypes characterized by mitochondrial dysfunction.

## DECLARATIONS

### Ethics approval and consent to participate and for publication

The study was approved by the Ethics Committee of the Healthcare System of the Autonomous Province of Bozen/Bolzano (approval number 102-2014 dated 26.11.2014 with extension from 19.02.2020) and of the CIUSSS de la Capitale Nationale. Brain donors provided written informed consent.

### Availability of data and material

All data generated during this study are included in this article and its supplementary information file.

### Authors’ contributions

CB, MPCR, MLP: Performed experiments, data analysis, wrote and reviewed the manuscript; GG, AZ, VSL, VC, VR, SL, CG, MRP, CH, PR, SK: performed experiments, data analysis, reviewed the manuscript; AAL: performed confocal imaging analyses, reviewed the manuscript; GB: performed electron microscopy analyses, reviewed the manuscript; GV: performed experiments and helped revise the manuscript; ML: performed experiments, data analysis, reviewed the manuscript; PPP, AAH: acquired funding, reviewed the manuscript; EZ, JS, VJ, MP, RB: supervised experiments and reviewed the manuscript; IP and ML: supervised the study, analyzed data, acquired funding, wrote and reviewed the manuscript. All authors read and approved the final manuscript.

## Acknowledgements

The authors acknowledge the Canadian Neurophotonics Platform of the CERVO Brain Research Centre for AAV production. We also appreciate the experimental support provided by Sara Meschini and Stephanie Marini, as well as valuable feedback from Rachel Morin-Pelchat, Jean-François Rivest, and Sandeep Sundara Rajan. The authors are grateful for brain donations and thank Marie-Josée Wallman for technical assistance. We also thank Dr. Anna Masato (University of Padova) for kindly providing the SH-SY5Y-tetON-αSyn cells stably expressing the tet-ON system for tetracycline-inducible expression of *SNCA*.

