## Supplementary Materials for "Modulation of SLP-2 expression protects against alpha-synuclein neuropathology by mitigating mitochondrial dysfunction"

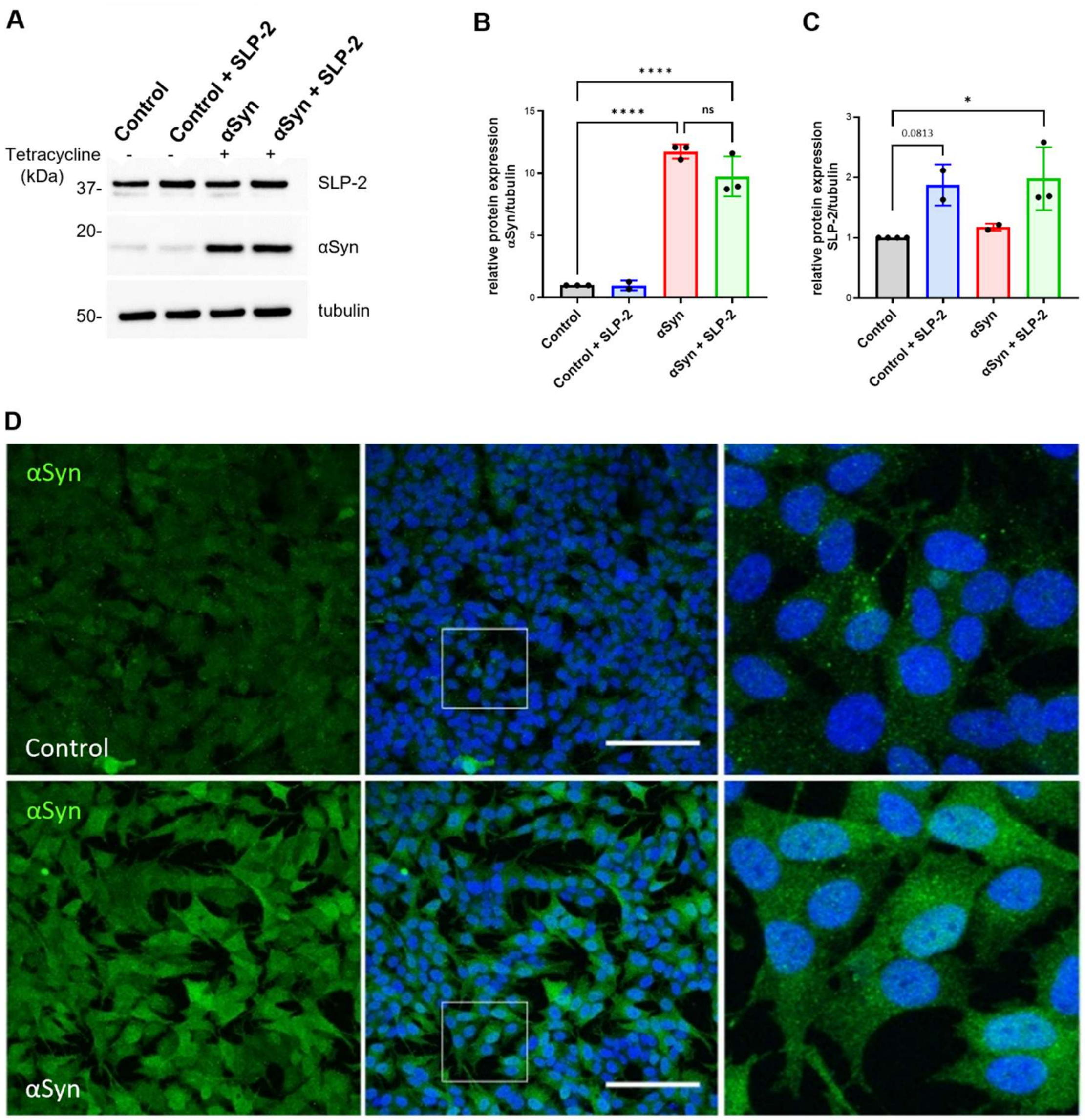


**Fig. S1. αSyn and SLP-2 protein levels after induction of wildtype αSyn and overexpression of SLP-2 in SH-SY5Y αSyn cells.** αSyn expression was induced in SH-SY5Y tetON-αSyn cells by addition of tetracycline. SH-SY5Y-tetON-αSyn were infected with a lentiviral construct carrying the SLP-2 coding sequence. (A) Whole cell lysates of uninduced (Control) and induced (αSyn) SH-SY5Y αSyn cells, before and after transduction with pER4-STOML2 lentivirus, were analyzed by Western blot with anti-SLP-2 and anti-αSyn antibodies. Tubulin served as loading control. (B, C) Densitometric analyses of αSyn and SLP-2 protein levels. Relative density values were normalized to the loading control tubulin. Molecular mass markers are in kilodaltons (kDa). Statistical differences were calculated by one-way ANOVA followed by Tukey's *post hoc* test to correct for multiple comparisons (n=3 independent experiments). Data is plotted as ± SEM. *p≤0.05; ****p≤0.0001. (D) Comparison of αSyn expression (green) in Control (uninduced) and αSyn-induced SH-SY5Y tetON-αSyn cells by immunofluorescence. Scale bar=100 µm.

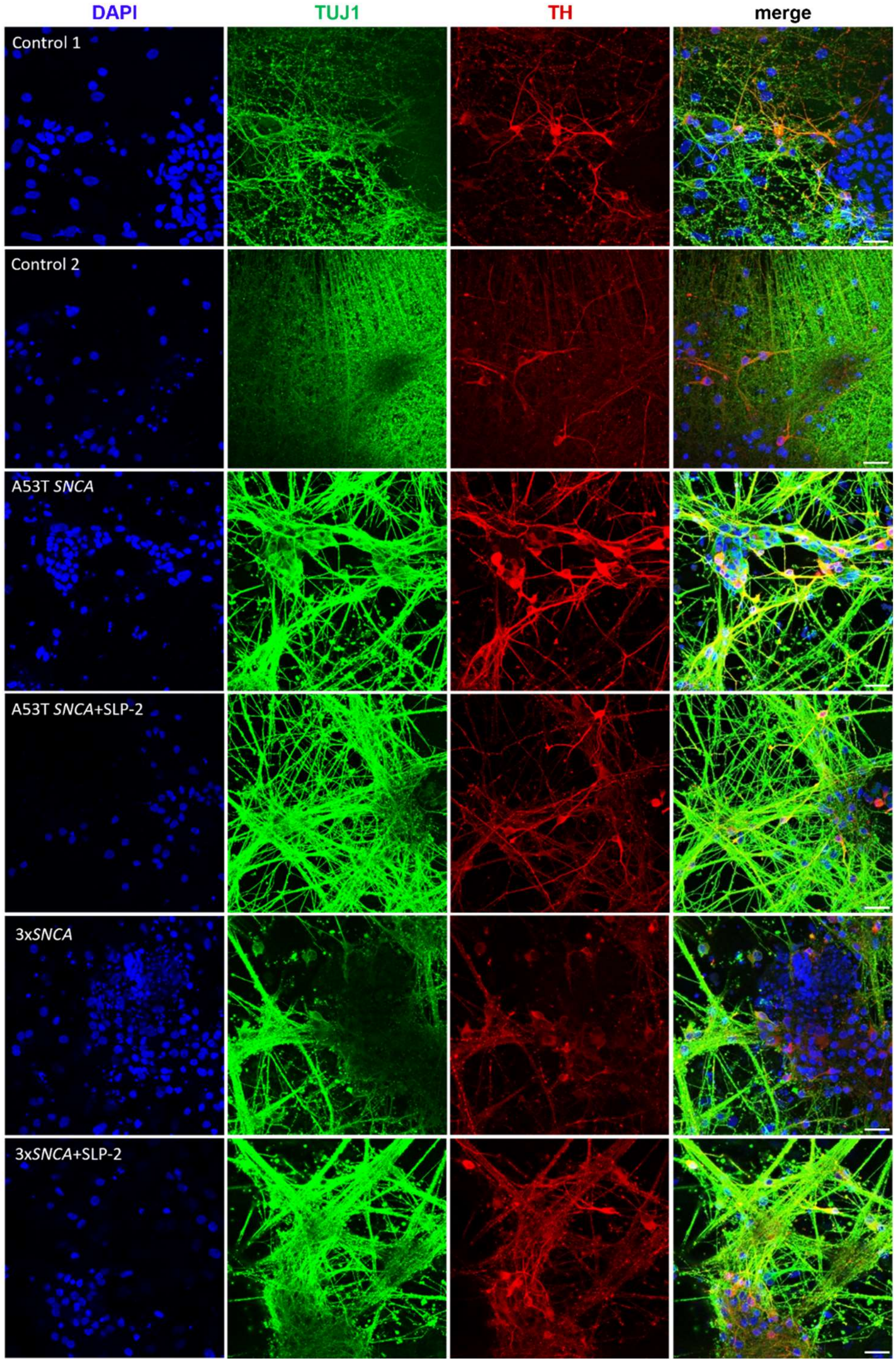


**Fig. S2. Immunocytochemical characterization of hiPSC-derived neurons with A53T or 3x*SNCA* mutations and healthy controls.** Immunofluorescence staining of neuronal cultures derived from two control lines and two *SNCA* mutation carriers (A53T *SNCA* and 3x*SNCA*) with and without SLP-2 overexpression (DIV 55), for neuron-specific TUJ1 (green), the DA marker TH (red), and nuclear DAPI (blue). Scale bar=25 µm.

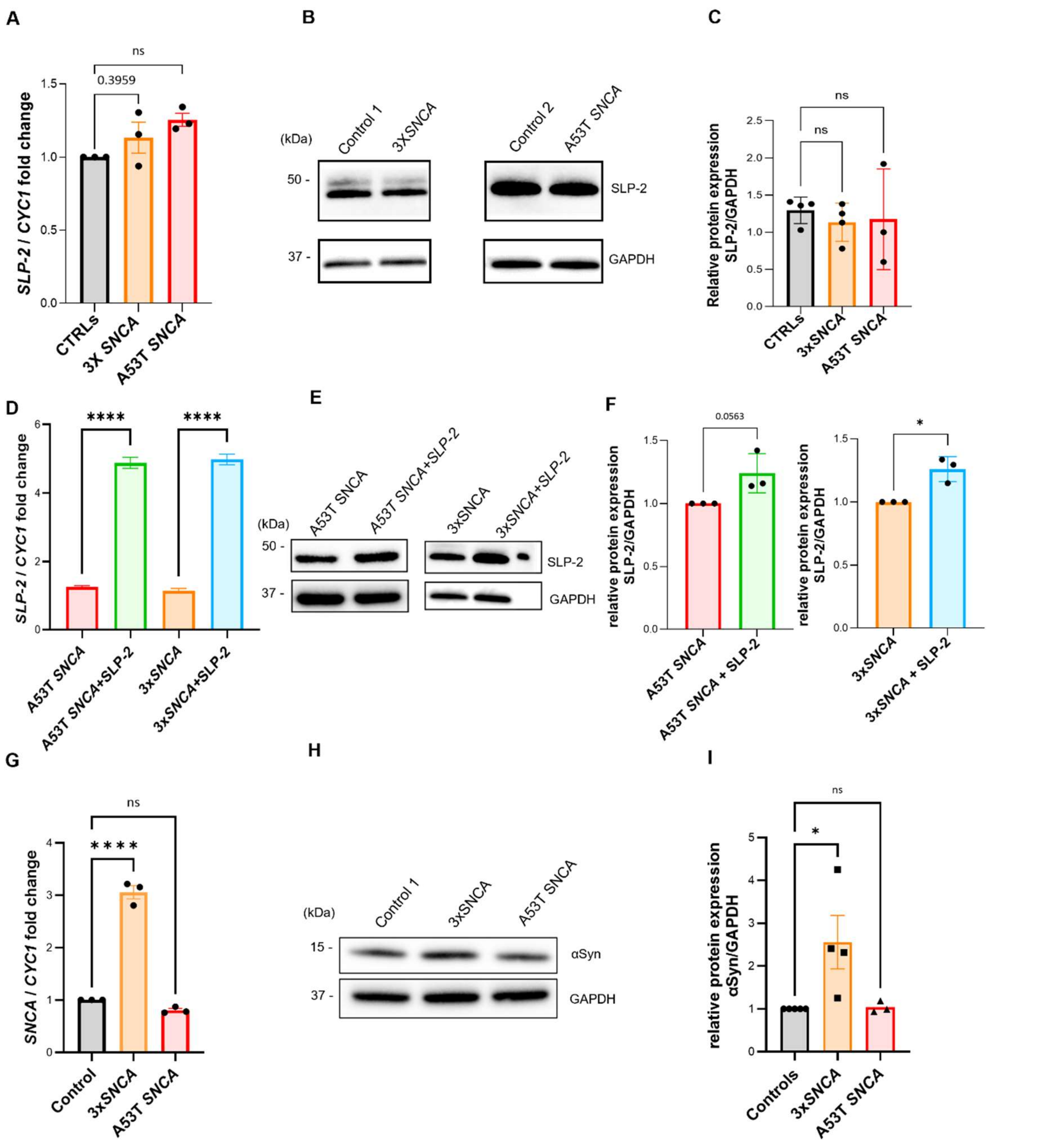


**Fig. S3. Characterization of hiPSC-derived neurons with A53T or 3x*SNCA* mutations and healthy controls**. (A) SLP-2 mRNA levels quantified by means of RT-PCR in hiPSC-derived neurons (DIV 45) in *SNCA* mutant lines compared to a control line. (B, C) Representative Western blot showing endogenous SLP-2 protein expression in hiPSC-derived neurons in *SNCA* mutant lines (3x*SNCA* and A53T *SNCA*) compared to control lines and the densitometric analysis. (D) Overexpression of *STOML2* mRNA levels quantified by means of RT-PCR in hiPSC-derived neurons in *SNCA* mutant lines compared to a control line. (E, F) Increase of SLP-2 levels was quantified by means of Western blot analysis in hiPSC-derived neurons in *SNCA* mutant lines compared to a control line and densitometric analyses. Viral SLP-2 transduction in the A53T *SNCA* and 3x*SNCA* lines induces 1.2-1.3-fold protein levels of SLP-2. (G) Total αSyn levels were analyzed by RT-PCR and (H-I) Western blot in the A53T *SNCA* and 3x*SNCA* lines. SLP-2 and total αSyn levels were analyzed with the indicated antibodies, and GAPDH was used as loading control. Molecular mass markers are in kilodaltons (kDa). n=3-4 independent experiments. Error bars represent the mean ± SEM. P-values were determined by a one-sided ANOVA test followed by Tukey's *post hoc* test to correct for multiple comparisons. *$p \leq 0.05$; ****$p \leq 0.0001$. ns=not significant.

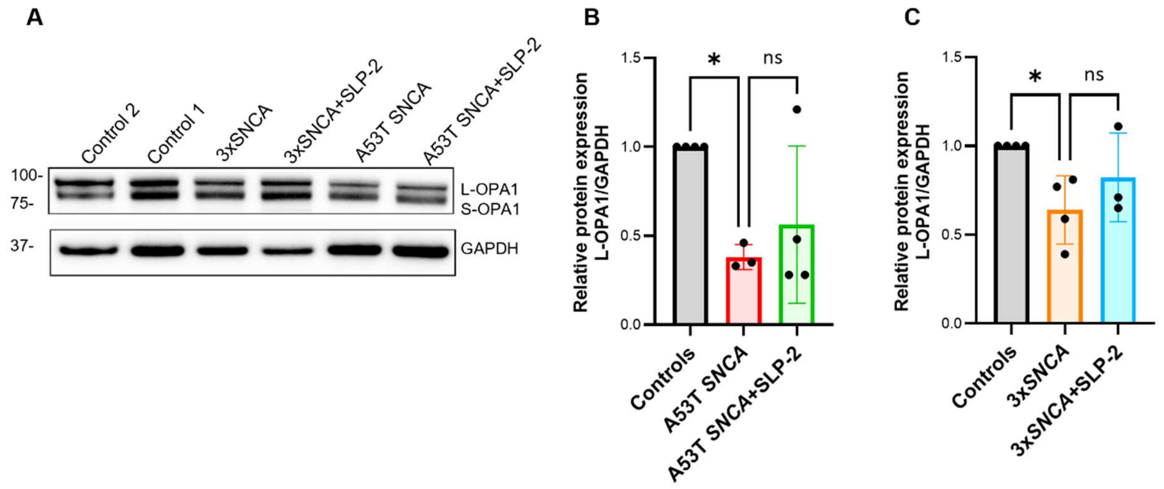


**Fig S4. SLP-2 overexpression rescues altered mitochondrial dynamics in hiPSC-derived neurons carrying *SNCA* mutations.** (A) Representative Western blot images of long (L-OPA1) and short (S-OPA1) isoforms of OPA1 in the hiPSC-derived neurons of controls, *SNCA* mutant lines and the *SNCA* mutant lines+SLP-2. GAPDH was used as a loading control. (B, C) Densitometric analyses of L-OPA1 in A53T *SNCA* and 3x*SNCA* mutant neurons, respectively. Relative density values were normalized to the loading control GAPDH. Molecular mass markers are in kilodaltons (kDa). n=3-4 independent experiments. Error bars represent the mean ± SEM. P-values were determined by a one-sided ANOVA test followed by Tukey's *post hoc* test to correct for multiple comparisons. *p≤0.05. ns=not significant.

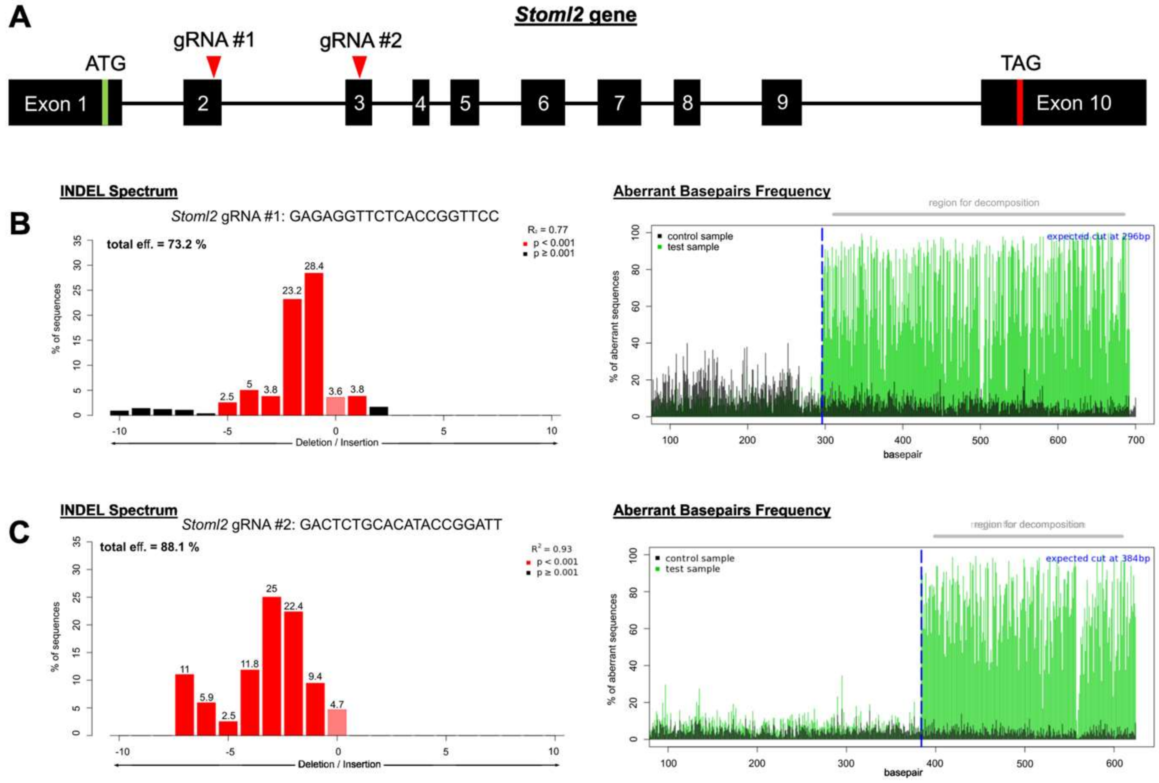


**FigS5. dgRNA against *Stoml2* gene efficiently cleaves their target exons.** Mouse fibroblasts NIH-3T3 were co-transfected with SpCas9 along with plasmids CAG-DIO-mCherry-NRN1-dgRNA-*Stoml2* or control with the plasmid CAG-DIO-mCherry-NRN1. Following selection with an antibiotic, the genomic DNA was extracted and sequenced by Sanger sequencing technology to assess the nucleotide Insertions and Deletions (INDELS) at the target sites. (A) Schematic of the mouse *Stoml2* gene. Exons are shown as black boxes. The start and stop codons' locations are shown as green and red stripes, respectively. The targets of the dgRNAs are shown with red arrows. (B) The spectrum of INDELS and aberrant basepair frequency passed the expected cut site measured by TIDE assay for the gRNA#1 and (C) gRNA#2.

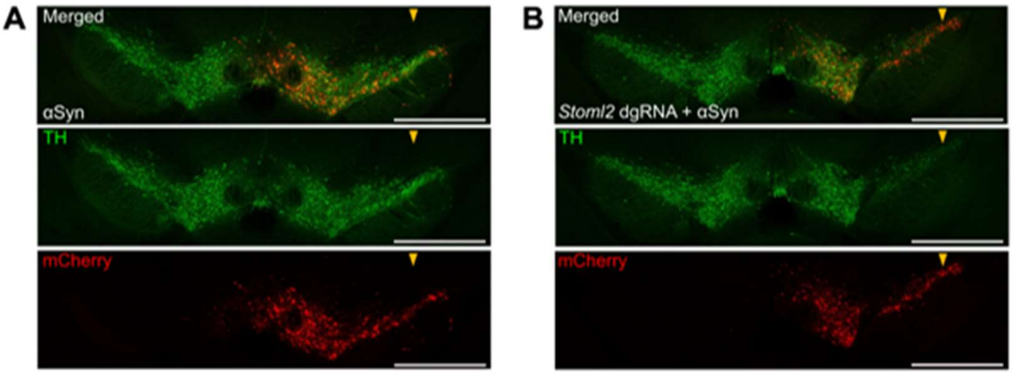


**FigS6. Supplementary panels for Fig. 10.** Representative images of mouse SNc coronal sections. The cell bodies of DA neurons were labelled for TH (green), and infection was confirmed with the presence of the mCherry reporter (red). The arrows indicate the side of the injection of the AAV. Scale bar = 1,000 µm.

**Table S1. Brain donors age, sex, *post-mortem* delay and cause of death.**

| Group | Identification | Age | Sex | Post-Mortem Delay (hr) | Cause of Death |
|---|---|---|---|---|---|
| **Parkinson's disease** | C-0002 | 78 | M | 2.5 | Heart/Kidney Failure |
| | C-0004 | 73 | F | 18 | Acute Pneumonia |
| | C-0018 | 68 | F | 14 | Bronchoaspiration |
| | C-0049 | 77 | M | 4 | Adenocarcinoma |
| | C-0052 | 79 | M | 7 | Aspiration Pneumonia |
| **Control** | H-42 | 71 | F | 5 | Lung Disorder |
| | H-154 | 73 | M | 20 | Subdural Hemorrhage |
| | H-172 | 72 | F | 20 | Coronary Arteriosclerosis |
| | H-486 | 75 | M | 12 | Rupture of the Aorta |
| | C-0040 | 93 | F | 24 | Cerebellar Hemorrhage |
| | C-0078 | 92 | F | 9.5 | Stroke and Epilepsy |
| | C-0089 | 55 | M | 15 | Glioblastoma |
| | C-0103 | 94 | M | 24 | Heart Failure |

**Table S2. Experimental resources (mouse strains, cell lines and virus strains).**

| MOUSE STRAINS | SOURCE | IDENTIFIER |
|---|---|---|
| C57BL/6 | Charles River | C57BL/6NCrl |
| $DAT^{IRES\text{-}Cre}$ | Jackson | 006660 |
| $Rosa26^{LSL\text{-}Cas9}$ | Jackson | 024857 |
| **CELL LINES** | **SOURCE** | **IDENTIFIER** |
| NIH-3T3 | ATCC | CRL-1658 |
| SH-SY5Y-tetON-αSyn | Dr. Elisa Greggio University of Padova) | N/A |
| hiPSC-802 | Institute for Biomedicine (Eurac Research) | N/A |
| EURACi014-A; hiPSC-1.1 | Institute for Biomedicine (Eurac Research) | FFF-091 (FFF-027 2014) |
| hiPSC-ND34391 | NINDS Human Cell and Data Repository | ND34391 |
| hiPSC-SFC 089-03-07-05A | StemBANCC consortium (https://cells.ebisc.org/STBCi033-B/) | SFC 089-03-07-05A |
| **VIRUS STRAINS** | **SOURCE** | **IDENTIFIER** |
| AAV2/9-hSyn-DIO-mCherry | Addgene | 50459 |
| AAV2/9-hSyn-DIO-SLP2-IRES-mCherry | this paper | N/A |
| AAV2/9-hSyn-SLP2-IRES-mCherry | this paper | N/A |
| AAV2D/J-hSyn-Con/Fon-hChR2(H134R)-EYFP | Addgene | 55645 |
| AAV2/9-CMVie-hSyn-αsynA53T | this paper | N/A |
| AAV2/9-CAG-DIO-mCherry-NRN1 | this paper | N/A |
| AAV2/9-CAG-DIO-mCherry-NRN1-U6-dgRNA-Stoml2 | this paper | N/A |
| AAV2/9-hSyn-mKate | this paper | N/A |
| AAV2/9-hSyn-SLP2-IRES-mCherry | this paper | N/A |
| Lentivirus/pER4-h*STOML2* | (*42*) | N/A |
